# Reversible repression of inducible genes by Polycomb Repressive Complex 2 and H3K27me3 in Drosophila melanogaster

**DOI:** 10.1101/2025.06.09.658596

**Authors:** Robert Streeck, Holly N. Stephenson, Alf Herzig

**Affiliations:** Department of Cellular Microbiology, Max Planck Institute for Infection Biology; Charitéplatz 1, Berlin 10117, Germany

## Abstract

The Polycomb system maintains cell fate decisions by establishing silenced chromatin at developmental regulators. This is mediated by histone H3 lysine 27 trimethylation (H3K27me3) deposited in highly modified chromatin domains by Polycomb repressive complex 2 (PRC2). Increasing evidence suggests that the Polycomb system has functions beyond developmental regulation but the underlying mechanisms remained unclear. Here, we show that large regions of the genome are moderately modified by H3K27me3 and transcriptionally repressed through PRC2. In contrast to genes in silenced chromatin, moderately modified genes remain inducible by physiological stimuli and the repressive function of PRC2 is balanced by activating inputs from the Trithorax system. This demonstrates a pervasive function of the Polycomb system in dynamically regulating homeostatic expression and inducibility of genes in differentiated cells.

**One-Sentence Summary:** Large regions of the genome are reversibly repressed by the Polycomb system, which prevents gene activation by low-level stimulation but maintains inducibility.

## Introduction

Studies in many model systems showed that the Polycomb system generates stably inherited domains of silenced chromatin in which genes remain inactive in the presence of their cognate transcriptional activators (*1-3*). In *Drosophila melanogaster*, Polycomb repressive complex 2 (PRC2) binds Polycomb response elements (PREs) at inactive genes during early embryogenesis, which leads to histone H3 lysine 27 trimethylation (H3K27me3) by the PRC2 subunit Enhancer of zeste (E(z)) (*1-4*). H3K27me3 is essential for transcriptional repression and together with continued recruitment of PRC2 at PREs ensures stable inheritance of silenced chromatin domains (*5-8*). At transcriptionally active genes, Trithorax group (TrxG) proteins prevent Polycomb dependent silencing. TrxG proteins are part of several alternative COMPASS complexes which are responsible for histone H3 lysine 4 methylation (H3K4me1, H3K4me3) (*9*). H3K4 methylation inhibits PRC2 activity and counteracts stable silencing (*3, 10*). Central to this paradigm is that transcriptional states of genes (ON or OFF) are read out during a brief window in development and OFF states become irreversible in differentiated cells. There is mounting evidence however, that the Polycomb system has functions in postembryonic development and in differentiated cells that may not fit this basic paradigm (*11-17*). Here, we use *Drosophila melanogsater* plasmatocytes, which are macrophage-like cells (*18*), to address these functions in an *in vivo* model.

## Results and Discussion

To identify Polycomb target genes we generated H3K27me3 ChIP-seq profiles, which we correlated to transcriptional activity (Fig 1A, Fig S1). Plotting average H3K27me3 signals per gene revealed a large population of genes modified above background, which was corroborated by standard MACS2 peak calling (p<0.05 Fig 1B). Known PRC2 target genes localized to the highly modified tail of this population, corresponding to genes overlapping highly stringent MACS2 peaks (p<10^−40^, Fig. 1B). Together this indicated that in addition to ‘canonical’, highly modified PRC2 targets, about 40% of plasmatocyte genes were moderately modified by PRC2. To better classify PRC2 target genes we developed the ClassFinder tool. ClassFinder uses multivariant binomial mixture models to fit ChIP-seq data to a given number of underlying populations by an expectation maximization algorithm. We found that a 3-population model best fit the data, separating highly modified (Pc-H), moderately modified (Pc-M), and non-modified genes (non-Pc) (Fig. 1B, Fig. S2). We then re-analysed H3K27me3 ChIP-seq data sets from diverse *Drosophila* cell types and found a comparable distribution of PRC2 target genes (Fig. S3).

**Fig. 1.**
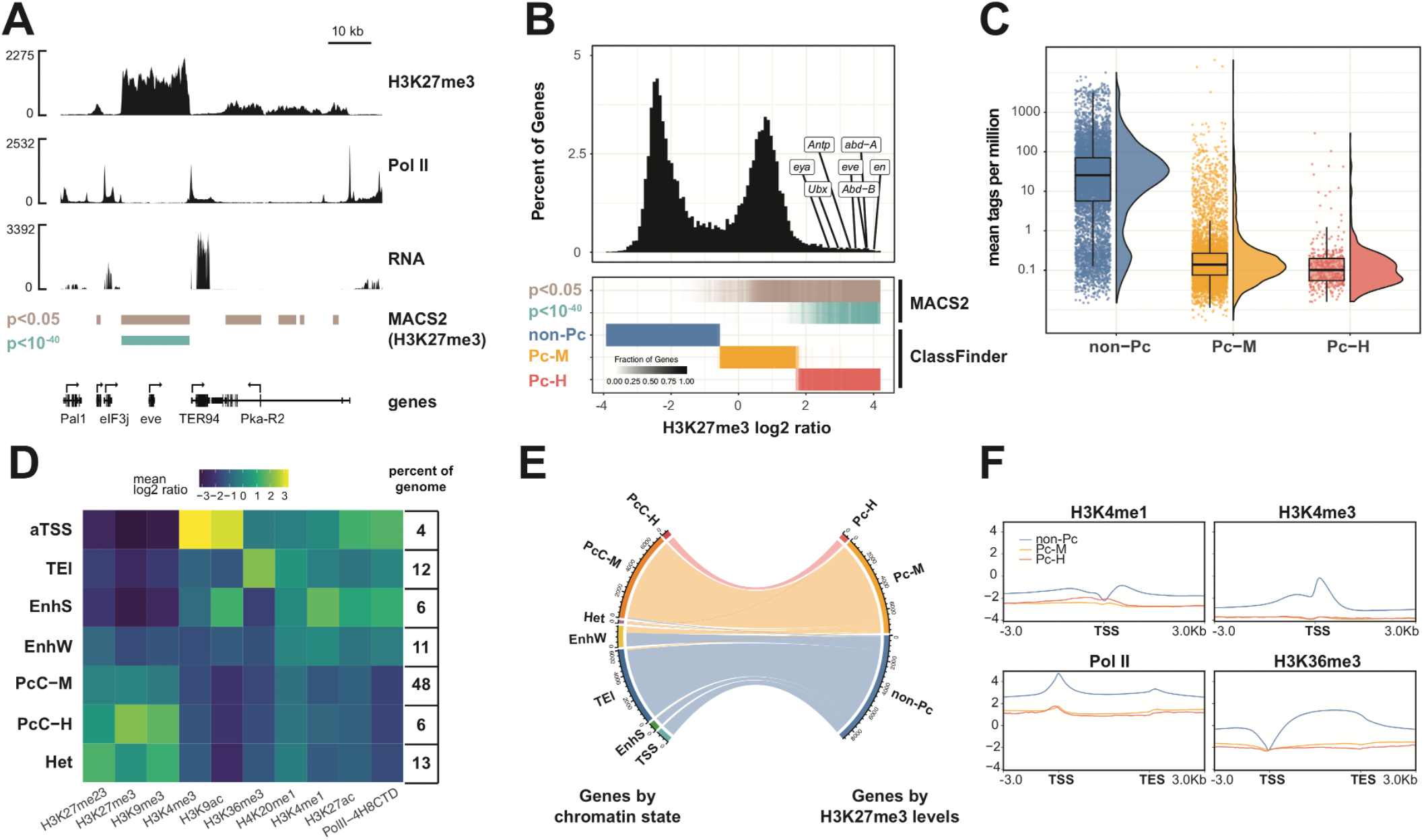
The Polycomb system targets a large group of moderately modified and repressed genes. (**A**) ChIP-seq tracks for H3K27me3 and RNA-polymerase II (Pol II) and RNA-seq track around the *even skipped* (*eve*) locus. H3K27me3 MACS2 peaks with highly significant cutoff (p<10^−40^, teal) and all peaks (p<0.05, brown) are shown. (**B**) Histogram of gene level H3K27me3 signal (top) and corresponding classification (bottom). H3K27me3 signal was determined as log2 ratio over all exons from H3K27me3 ChIP-seq data. Canonical PRC2 targets are marked by labels. (**C**) Relative expression of genes determined by mean gene level tags per million from RNA-seq. Resulting distributions are shown for ClassFinder gene sets (non-Pc, Pc-M, Pc-H). (**D**) ClassFinder fit of ChIP-seq data sets (below heatmap) over 200 bp bins. Heatmap shows abundance of ChIP-seq features across a 7-state model with genome coverage indicated in percent (aTSS, active transcriptional start sites; TEl, transcriptional elongation; EnhS/EnhW, enhancer strong/weak; PcC-M/PcC-H, Polycomb moderate/high; Het, constitutive heterochromatin). (**E**) Mapping of genes between H3K27me3 gene states and modification states. (**F**) Average distribution of ChIP-seq signals for individual gene states (non-Pc, Pc-M, Pc-H). H3K4me3, H3K27ac and H3K4me1 and were mapped around the transcriptional start site (TSS); Pol II along the gene body.

Since transcriptional inactivity is a key feature of canonical PRC2 target genes, we asked whether moderately modified PRC2 targets were silent as well. Median expression levels of Pc-M and Pc-H genes by RNA-seq were similarly low, but the distribution showed a tail of weakly expressed Pc-M genes (Fig. 1C). This weak expression could be explained by transcriptional diversity within the plasmatocyte population. We therefore assessed Pc-M genes in existing expression data of plasmatocyte subpopulations that were identified by single cell RNA-seq (*19*). Pc-M genes were either not detected or weakly expressed in all of the individually annotated clusters, which resembled their expression profile in the whole population (Supp Fig. 4). For in-depth comparison of canonical and moderately modified PRC2 target genes we generated additional ChIP-seq data sets (Fig. S5, S1). Similar to the Pc-M/Pc-H gene classification we trained the ClassFinder algorithm to assign genome bins to 7 modification states. This model predicted states that matched known chromatin states (*20*) and two Polycomb chromatin states (PcC-M, PcC-H) that were primarily discriminated by repressive chromatin modifications such as H3K27me3, H3K9me3 and H3K27me2/3 (Fig. 1D, Figs. S6-7). Genome coverage of PcC-H chromatin (6%) closely resembled that of Polycomb chromatin in previous models (*20*). Pc-M and Pc-H genes (defined by H3K27me3 alone) almost exclusively mapped to the respective modification states, indicating that H3K27me3 is a key characteristic for defining these groups (Fig 1E). To compare both groups with respect to chromatin marks found at active promotors and enhancers we analyzed gene level modification profiles. Relative to non-modified genes, Pc-M and Pc-H genes were similarly depleted in H3K4me1, H3K4me3, H3K27ac and active Pol II (Fig. 1F). Pc-M genes as a group, therefore show no bivalency for active and repressive chromatin marks. Nevertheless, we found a gradual increase in active chromatin marks within the group of Pc-M genes that correlated with a decrease in H3K27me3 levels (Fig. S8) which is consistent with a subset of Pc-M genes being weakly expressed (Fig 1C).

Taken together, the data suggested that Pc-M genes could be at the brink of escaping PRC2 dependent repression. Therefore, we asked whether increasing transcriptional stimulation could induce their expression. We stimulated the IMD and JAK/STAT signaling pathways by GAL4/UAS dependent overexpression of the IMD pathway receptor PGRP-LC or the JAK kinase hopscotch in plasmatocytes (*21, 22*). In order to pulse pathway activation, we used the plasmatocyte-specific and mifepristone-dependent Hml-GAL4^S^ driver line (*23*) (Fig. 2A). After a 24 h mifepristone pulse we detected a strong response to the hormone, similar to previous observations (*24*) (Fig. S9). However, within IMD and JAK/STAT specific target genes (Fig. 2A) and within all up-regulated genes (Fig. S9), Pc-M genes were significantly enriched. This suggested that Pc-M genes were inducible. To physiologically stimulate plasmatocytes we infected larvae with a mix of gram-positive and gram-negative bacteria. We found a transient transcriptional response that returned close to baseline after 18 h (Fig. 2B, Fig. S10). Immune induced genes at all timepoints were enriched for Pc-M genes, showing that Pc-M genes are part of the physiological response to infection. Next, we addressed whether at the level of individual genes up-regulation of Pc-M genes was associated with a loss of H3K27me3 and a gain of H3K27ac. We generated H3K27me3 and H3K27ac ChIP-seq profiles of plasmatocytes 6 h post-infection and normalized these to profiles of unchallenged plasmatocytes (Fig S11). Pc-M genes that were induced 3 h and 6 h post-infection lost the repressive H3K27me3 modification and gained H3K27ac in return (Fig 2C, Fig S12). We also found a correlation between transcriptional induction and the loss of H3K27me3 at Pc-M genes (Fig 2D, Fig S12). This indicated a dynamic and quantitative relationship between the transcriptional output of genes and their H3K27me3 modification level.

**Fig. 2.**
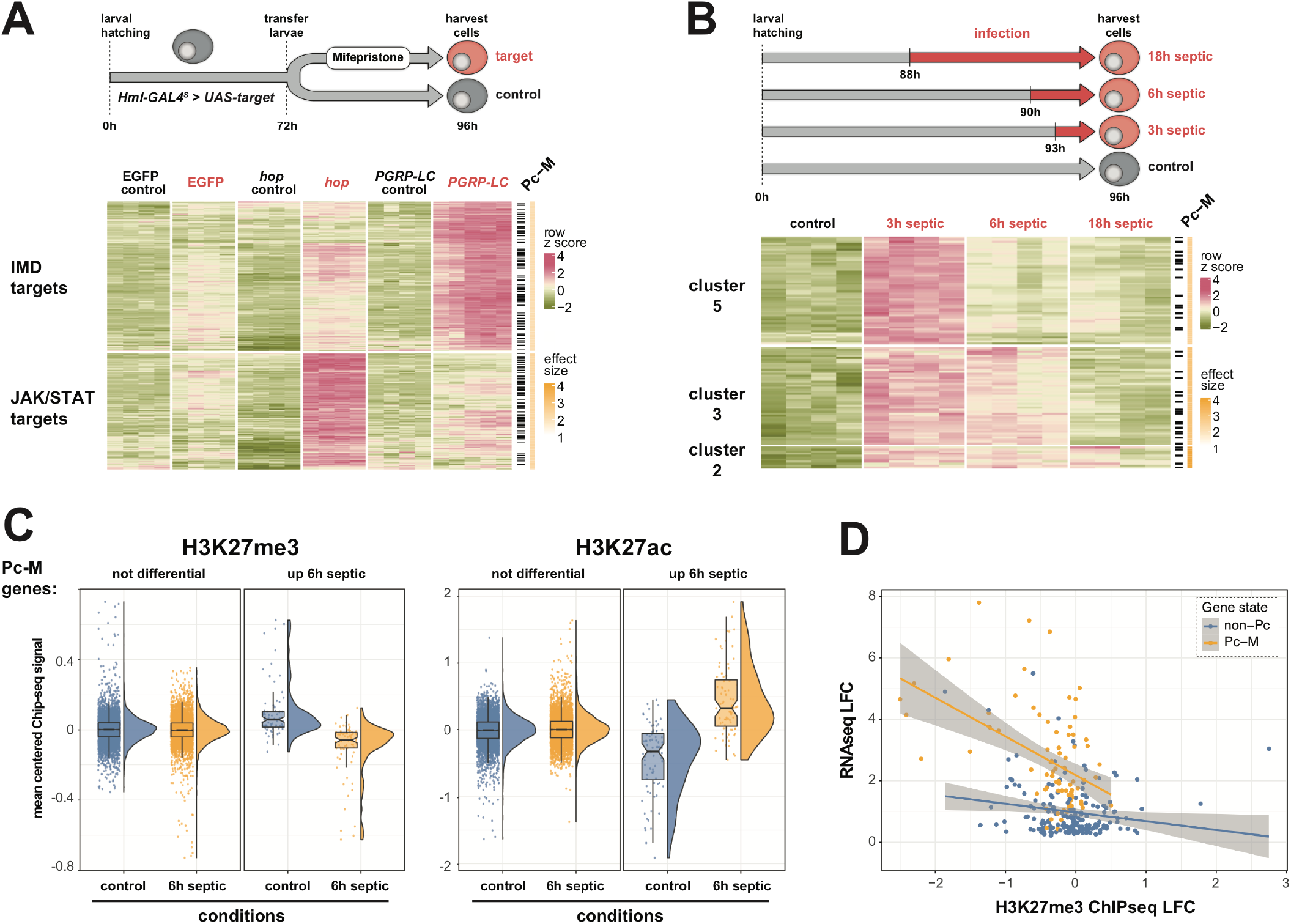
Moderately modified Polycomb targets are inducible and moderate H3K27me3 levels are reversible. (**A)** Plasmatocyte specific expression of *EGFP, hopscotch* (*hop*) and *PGRP-LC* was induced by mifepristone and compared to non-induced conditions by RNA-seq. Heatmap shows normalized row z-scores for up-regulated *hop* and *PGRP-LC* target genes (full heatmap fig. S9). Pc-M genes are indicated by black bars and effect size of Pc-M gene enrichment by yellow color scale for each cluster. (**B**) Plasmatocytes were isolated 3 h, 6 h or 18 h after septic injury from 96 h old third instar larvae. Heatmap shows normalized row z-scores for genes up-regulated compared to control plasmatocytes by RNA-seq (full heatmap fig. S10). Pc-M genes are indicated by black bars and effect size of Pc-M gene enrichment by yellow color scale for each cluster. (**C**) Differential modification of plasmatocyte Pc-M genes 6 h after septic injury determined by H3K27me3 or H3K27ac ChIP-seq. Pc-M genes were grouped into genes either up-regulated or not differentially expressed 6 h after septic injury. For each gene in these groups mean centered ChIP-seq signals were determined in control plasmatocytes (blue) and in plasmatocytes 6 h after septic injury (yellow). (**D**) For genes up-regulated at 6 h post septic injury, ChIP-seq mean log2 fold change (from quantile normalized data) is plotted against RNA-seq log2 fold change. Pc-M genes are marked in yellow and non-Pc genes in blue. Lines show linear models with 95% confidence interval in grey. For Pc-M genes there is a statistically significant correlation (r = -0.48, p = 7.8*10^−6^).

We showed that moderately modified PRC2 targets were not irreversibly silenced, but in fact inducible when homeostatic conditions were perturbed by transcriptional stimulation. Next, we addressed whether PRC2 regulates the expression of Pc-M genes under homeostatic conditions. Canonical PRC2 targets that are silenced against homeostatic activation are up-regulated in differentiated cells upon elimination of PRC2 function or replacing histone H3 with modification-resistant histone H3K27R (*7, 8, 25*). We generated plasmatocytes mutant for the PRC2 subunits E(z) and Supressor of zeste 12 (Su(z)12), or with H3K27R replacement using the flp/FRT system (*26*) and compared them to control cells by RNA-seq (Fig. 3). Clustering of affected genes showed that canonical PRC2 targets (Pc-H) and moderately modified Pc-M genes were significantly enriched within up-regulated genes (Fig. 3, Fig. S13). We then compared the gene-sets repressed by PRC2 with the gene-sets up-regulated upon transcriptional stimulation (Fig. S14). We found a significant overlap between both gene sets, showing that PRC2 repression extends to inducible genes. This included ∼50% of immune induced Pc-M genes indicating that the Polycomb system suppresses immune-related responses to continuous low-level stimulation in plasmatocytes.

**Fig. 3.**
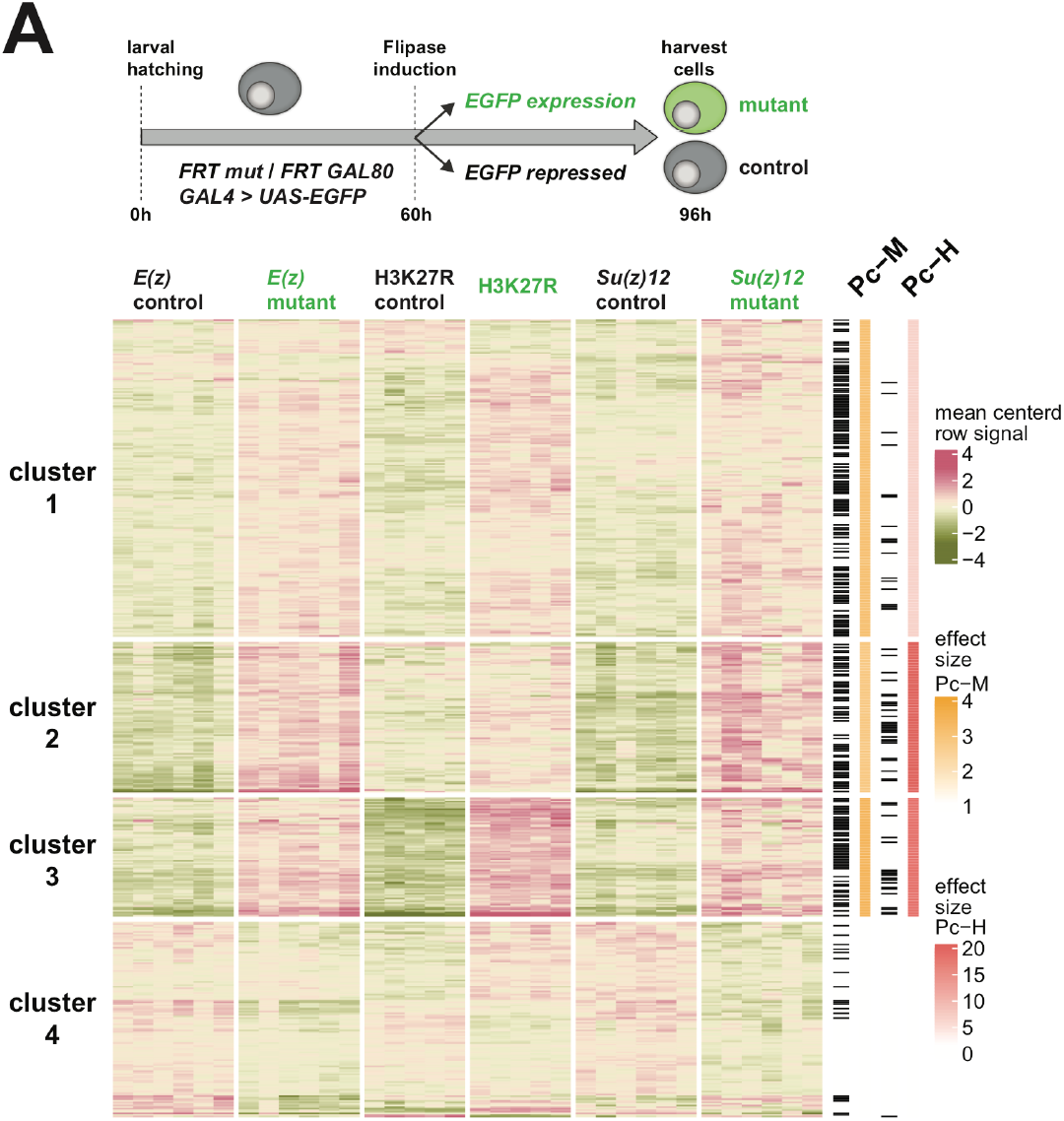
Moderately modified Polycomb targets are repressed by PRC2 and H3K27me3. (**A**) *E(z), Su(z)12* and H3K27R mutant plasmatocytes were induced by flp/FRT mediated recombination and compared to non-mutant sibling cells by RNA-seq. Heatmap shows normalized mean centered signals of differentially expressed genes. Pc-M and Pc-H genes are indicated by black bars and effect sizes of gene enrichment by yellow color scale (Pc-M) or red color scale (Pc-H) for each cluster. Pc-M and Pc-H genes were significantly enriched amongst up-regulated genes (clusters 1-3), but not within down-regulated genes (cluster 4).

We showed that the level of H3K27me3 at Pc-M genes is dynamically reduced by transcriptional activation (Fig. 2). At canonical PRC2 targets, TrxG proteins mediate a comparable effect of transcriptional activation in early development (*27, 28*). Key TrxG proteins are the H3K4 methyltransferases TRR, TRX and dSET1 that are part of alternative COMPASS/COMPASS-like complexes (*9*). TRR/COMPASS is responsible for H3K4 monomethylation at most enhancers and also contains the H3K27me3 demethylase UTX (*29, 30*). TRX/COMPASS counters PRC2 at canonical PRC2 targets by H3K4 trimethylation in early development (*31, 32*); whereas dSET1/COMPASS is responsible for H3K4 trimethylation at most actively transcribed genes (*30, 33*). We used plasmatocyte-specific RNAi knockdown to limit the expression of *trr, Utx, trx*, and *dSet1* and assayed the effect on Pc-M genes (Fig 4A, Fig. S15). Depletion of all four factors had a negative effect on gene expression in resting plasmatocytes (Fig. S16). Clustering of down-regulated genes showed that Pc-M genes were most prominently enriched in gene-sets that were co-dependent on TRX/COMPASS and TRR/COMPASS (Fig 4A). We found no significant enrichment of Pc-M genes in clusters that were dependent on dSET1/COMPASS, consistent with the idea that this complex preferentially acts on highly transcribed genes. Interestingly, we found genes that were co-dependent on TRX and UTX but not on TRR, which challenges the strict association of UTX with TRR/COMPASS (Fig. 4A). Genes down-regulated after *trx, trr* and *Utx* knockdown and genes up-regulated upon PRC2 inactivation showed significant overlap (Fig. 4B). Together with our previous observations, this indicated that the antagonistic function of the Polycomb and Trithorax systems extends to the regulation of weakly expressed and inducible genes in differentiated cells and is not limited to the establishment of stable ON/OFF states in early development.

**Fig. 4.**
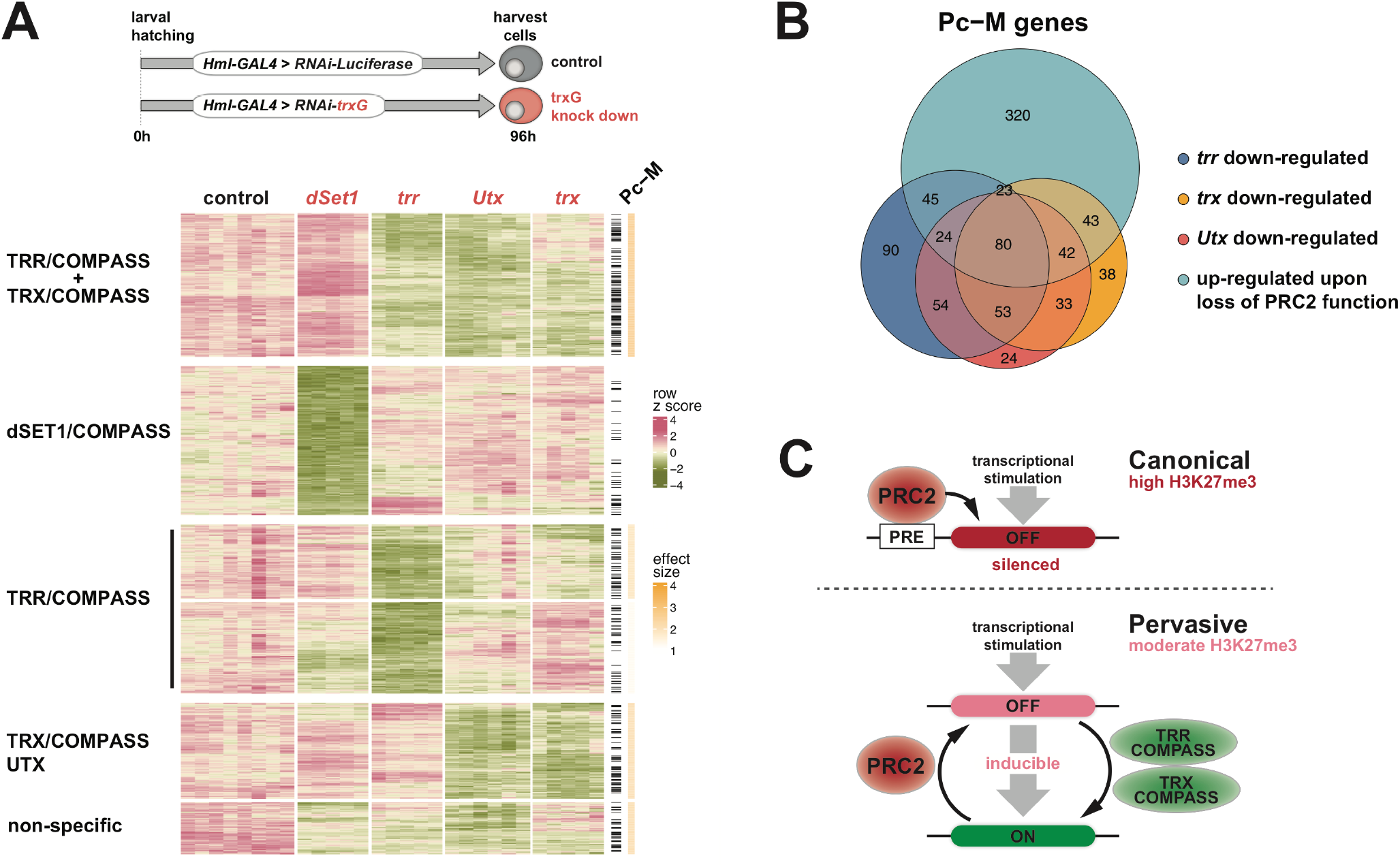
Repression is balanced by the Trithorax system at moderately modified target genes. (**A**) *dSet1, trr, Utx, trx* and *Luciferase* (control) were knocked down and compared by RNA-seq. Heatmap shows k-means clustering of normalized row z-scores. This identified genes primarily dependent on dSET1/COMPASS, TRR/COMPASS, TRX/COMPASS, genes dependent on TRX/COMPASS and UTX or no specific complex (non-specific). Pc-M genes are indicated by black bars and effect size of Pc-M gene enrichment by yellow color scale for each cluster. (**B**) Venn diagram of Pc-M genes down-regulated in *trr* (blue), *trx* (yellow) or *Utx* (red) knockdowns overlapping genes up-regulated in any PRC2/H3K27me3 mutant condition (see Fig. 4, teal). Significance of overlap by hypergeometric test: p < 10^−173^, effect size = 5.9. (**C**) Model for PRC2 function at canonical Polycomb targets compared to pervasive regulation of genes. See text for details.

In summary, we propose that the Polycomb system has a pervasive role regulating the homeostatic expression of genes beyond its established role in chromatin silencing (Fig 4C). An important aspect of our model is that transcriptional stimulation can reverse H3K27me3 dependent repression at moderately modified genes but not at canonical PRC2 targets. Inherited silencing of canonical targets requires persistent recruitment of PRC2 to PRE sites (*5, 6*). We mapped several sets of experimentally verified PREs to our chromatin state model and found them enriched in highly modified Polycomb chromatin (Fig. S17). This suggests that the difference in H3K27me3 levels between highly and moderately modified Polycomb chromatin could primarily reflect PRE-targeted versus pervasive recruitment of PRC2. Interestingly, a PRE-regulated transgene became inducible in a pattern reflecting the strength of endogenous transcriptional stimulation when the level of H3K27me3 was reduced (*5*). This suggests that the principal difference between silenced and reversibly repressed Polycomb chromatin is the degree of H3K27me3 modification, which at canonical PRC2 targets is not reversible by physiologically relevant stimulation. Mathematical models suggested that without PRE-targeted binding of PRC2 balanced inputs from the Polycomb and Trithorax systems create bistable chromatin that switches between ON and OFF states (*34, 35*). We found that weakly expressed genes are under balanced inputs from both systems, and chromatin bistability could explain why we found transcription in the apparent presence of H3K27me3. However, we did not detect a general increase in active chromatin marks or Pol II binding in the group of moderately modified Polycomb targets, which suggests that the majority of these genes are in an OFF state. The reversible mode of Polycomb dependent repression that we describe here extends the scope of the Polycomb-Trithorax systems and may help to understand their role in processes beyond the traditional developmental paradigm (*11, 13-15*)

## Materials and Methods

### Bacterial cultures for infections

*Escherichia coli* strain DH5*α* and *Staphylococcus aureus* strain 8325-4 (*36, 37*) were plated from frozen glycerol stocks on LB agar plates and grown overnight at 37°C. Plates were stored at 4°C for up to one week. To harvest the bacteria and generate bacterial pellets for septic injury, bacterial overnight cultures of *E.coli* and *S.aureus* were pooled at a 1:1 OD_600_ ratio and centrifuged at 4000 g for 10 minutes at room temperature.

### Fly husbandry

Flies were raised and maintained on standard cornmeal food at 25°C or room temperature. For controlled density cultures adult fly collections were left to lay eggs on apple juice agar plates with yeast paste overnight at 25°C. After 24 h hatched larvae were washed off the plate and further larvae were allowed to hatch for 2 h. Larvae were picked off the plate and transferred into 36 mm food tubes with yeast paste at a density of 120 larvae per tube and raised at 25°C (referred to as ‘controlled density’ from here on). Larvae reached third instar wandering stage 96 h after hatching, at which stage plasmatocytes were extracted. Genotypes of all stocks used are described below.

### Switch GAL4 (mifepristone induced expression)

Virgin females from stock *P*{*GS.HmlΔ-GAL4*} were crossed to males from UAS-lines (*P*{*w[+mC]=UAS-PGRP-LC.a*}*3, P*{*w[+mC]=UAS-hop.H*}*2, P*{*UAS-2xEGFP*}). From these crosses, first instar larvae were collected 0-4 h after hatching and grown at controlled density on standard food for 72 h at 25°C, then transferred to new vials with standard food or standard food supplemented with 10 µM mifepristone (10 mg/ml stock solution in EtOH). Plasmatocytes were harvested after 24 h incubation at 25°C by bleeding.

### RNAi knockdowns

Virgin females from stock w[*];;*P*{*Hml[e9-P2A]-GAL4*}*attP2* (*38*) were collected and crossed to males from TRiP RNAi lines obtained from Bloomington stock center (*dSet1*: BL 33704; *trr*: BL 6916; *Utx*: BL 34076; *trx*: BL 33703; *Luciferase*: 31603). From these crosses, larvae were picked as above.

### Septic injury

Sharpened tungsten needles were generated by electrolytically removing material from the tip of 0.25 mm tungsten wire. A piece of tungsten wire was connected to a 6V AC power supply and dipped into a 10% solution of NaOH in which the counter-electrode was inserted. Periodically, the wire was inspected for the progress of sharpening.

Sharpened tungsten needles were used to perform septic injury. Larvae were taken from food tubes by washing out all larvae with a 20% sucrose solution. Larvae were quickly washed in PBS, then dried on paper towels and transferred to a rubber dish. Larvae were then pricked with the tungsten needle dipped into the bacterial pellet on the side of one of the most posterior segments. Larvae were then transferred back into normal food tubes with 3 drops of PBS (to prevent drying) and grown at 25°C for the indicated time. For infection time courses plasmatocytes were always extracted around the same time of day, 92 h post hatching. Injuries were performed 3 h, 6 h or 18 h prior to this.

### Genetic elimination of PRC2 activity

Plasmatocytes were extracted from larvae in third instar wandering stage that were heat shocked twice; once between 24 h - 48 h after egg laying and once between 48 h – 72 h after egg laying, to induce expression of *flipase* for FRT mediated somatic recombination. Mutant and control plasmatocytes were isolated based on GFP expression by FACS from larvae:

1. *E(z*), mutant (GFP positive) and control (GFP negative): *P*{*ry[+t7.2]=hsFLP*}*1, P*{*w[+mC]=tubP-GAL4*}*1, P*{*w[+mC]=UAS-GFP.T:Myc.T:nls2*}*1*,
*y[1] w[*] / w[*];*
*E(z)[731], P*{*1xFRT.G*}*2A / P*{*w[+mC]=tubP-GAL80*}*LL9 P*{*w[+mW.hs]=FRT(w[hs]*)}*2A*
2. *Su(z)12*, mutant (GFP positive) and control (GFP negative), from larvae: *P*{*ry[+t7.2]=hsFLP*}*1, P*{*w[+mC]=tubP-GAL4*}*1, P*{*w[+mC]=UAS-GFP.T:Myc.T:nls2*}*1*,
*y[1] w[*]* / *w[*]*;
*Su(z)12[4], P*{*w[+mW.hs]=FRT(w[hs]*)}*2A, e[1] / P*{*w[+mC]=tubP-GAL80*}*LL9 P*{*w[+mW.hs]=FRT(w[hs]*)}*2A*
3. H3K27R, mutant (GFP negative): *P*{*ry[+t7.2]=hsFLP*}*1, y[1] w[*] / w[*]; Df(2L)HisC, P*{*neoFRT*}*40A /*
*P*{*w[+mC]=Ubi-GFP(S65T)nls*}*2L, P*{*ry[+t7.2]=ftz-lacC*}*USC1; M*{*3xHisGU.K27R*}*ZH-86Fb, Pbac*{*3xHisGU. K27R*}*VK00033 / M*{*3xHisGU. K27R*}*ZH-86Fb, Pbac*{*3xHisGU*.
*K27R*}*VK00033*
4. H3K27R, control (GFP negative): *P*{*ry[+t7.2]=hsFLP*}*1, y[1] w[*] / w[*]; Df(2L)HisC, P*{*neoFRT*}*40A /*
*P*{*w[+mC]=Ubi-GFP(S65T)nls*}*2L, P*{*ry[+t7.2]=ftz-lacC*}*USC1; M*{*3xHisGU.wt*}*ZH-86Fb*, *Pbac*{*3xHisGU.wt*}*VK00033 / M*{*3xHisGU.wt*}*ZH-86Fb, Pbac*{*3xHisGU.wt*}*VK00033*

Larvae were identified by phenotypic markers from crosses set up using the stocks listed in materials section. The crosses were left to lay eggs for 4 h using 20 female and 10 male flies to control larval density.

### Plasmatocyte extraction

For plasmatocyte extraction, treated or untreated third instar wandering stage larvae were collected from food tubes and washed in PBS. At the same time Schneider’s medium with 10% FCS and 10 mM N-Acetyl-L-Cysteine (pH 7.0) was prepared in a well of a 24-well tissue culture plate. Larvae were dried on paper towels and transferred to the tissue culture well with medium. Then they were bled by holding the head with forceps and with another pair of forceps ripping open the cuticula along the body of the larvae without disrupting the gut. 80 larvae were bled for each sample. The larval carcasses were removed and the plasmatocytes were allowed to attach for 10-15 minutes. The plasmatocytes were then washed 4 times with PBS.

### Plasmatocyte isolation and FACS sorting

For FACS sorting plasmatocytes were extracted as described above, with the exception that Nunclon Sphera ultra-low attachment dishes were used for bleeding instead. Cells were not washed with PBS but instead passed through 70 µm cell FlowMi cell strainers. DAPI was added to a final concentration of 1 µg/ml to mark dead cells.

Sorting of cells for RNA isolation was performed using a BD FACSAria II cell sorter, sorting for GFP using a 488 nm laser for excitation and a 530/30 filter for emission and sorting for DAPI negative cells using a 405 nm laser for excitation and a 450/40 filter for emission. The resulting cells were collected, spun down (2000 g, 4°C, 5 minutes) and lysed either in 500 µl TRIzol (H3K27R) or 94.5 µl RNAdvance lysis buffer (*Su(z)12* and *E(z)*). TRIzol samples were stored at -80°C, RNAdvance samples were isolated immediately.

### RNA isolation by TRIzol

Frozen TRIzol samples were thawed at room temperature, adjusted to 900 µl with TRIzol and moved to fresh prespun phase lock heavy tubes. 250 µl chloroform was added to each sample, mixed thoroughly and centrifuged (12000 g, room temperature, 15 minutes). The upper aqueous phase was then moved to a fresh DNA LoBind tube, making sure not to disrupt the interphase or phenol phase, and mixed by inverting with 550 µl isopropanol and 1 µl glycogen (20 mg/ml, RNAse free). Samples were incubated for 30 minutes at -20°C, then centrifuged (16000 g, 4°C, 10 minutes), and the supernatant was removed carefully without disrupting the pellet. The pellet was resuspended in 100 µl ultra-pure water with 300 mM sodium acetate and 1 µl glycogen (20 mg/ml, RNAse free). Next, 300 µl EtOH was added and the sample was incubated for 20 minutes at -20°C, then centrifuged (16000 g, 4°C, 10 minutes) and the supernatant was discarded. The pellet was washed two times by adding 1 ml of 70% EtOH (prepared with ultra-pure water), each time spinning down the pellet (16000 g, 4°C, 3 minutes). Afterwards all supernatant was drained and the pellet was dried until no liquid was visible. The pellet was then resuspended in 15 µl of ultra-pure water and stored at - 80°C.

### RNA isolation by RNAdvance

For direct RNA extraction, plasmatocytes were extracted as described above, then all supernatant was aspirated, they were lysed in 94.5 µl RNAdvance lysis buffer. RNA was isolated from freshly lysed plasmatocytes samples using the RNAdvance cell v2 kit following manufacturer’s instructions using a DynaMag-2 magnetic rack. DNase digestion was performed as suggested in the manual using Ambion DNase. Resulting RNA samples were eluted in 20 µl ultra-pure water and stored at -80°C.

### RNA quality control

RNA concentration of RNA samples for library preparation was measured using the Qubit RNA high sensitivity kit and the Qubit 2.0 fluorometer using 2 µl of RNA sample according to manufacturer instructions. 1 µl of RNA was then diluted to 1 ng/µl and run on an Agilent bioanalyzer RNA Pico. If samples showed the characteristic ribosomal RNA peaks without signs of degradation, they were used for subsequent library preparation.

### RNA library preparation

Libraries from FACS isolated plasmatocytes (H3K27 mutant, control) and from plasmatocytes of RNAi knockdowns were generated by the Max Planck Genome Centre (Cologne). Libraries from Switch GAL4 experiments were generated by Eurofins MWG. Libraries from FACS isolated plasmatocytes (*E(z*) and *Su(z)12* mutant and controls) and septic injury were generated using the NEBNext Ultra II Directional RNA Library Prep with Sample Purification Beads, the NEBNext Poly(A) mRNA Magnetic Isolation Module and the NEBNext Multiplex Oligos for Illumina sets according to manufacturer’s instruction. Adaptor concentrations were adjusted according to manual instructs. The number of PCR cycles was adjusted depending on starting RNA amount. The final libraries were eluted into 12 µl ultra-pure water and stored at -20°C.

### Plasmatocyte isolation and crosslinking for ChIP

Plasmatocyte were extracted from Oregon-R larvae 94 h after larval hatching (untreated or 6 h septic injury) as described above. After washing, plasmatocytes were fixed for 10 minutes at room temperature with 1,8% Formaldehyde in 1 ml PBS. Fixation was quenched for 3 minutes using 100 µl 2 M glycine. Cells were then scraped off the dish using a cell scraper and transferred to a 1.5 ml tube on ice. After a 5 minute incubation on ice, plasmatocytes were centrifuged (3000 g, 4°C, 5 minutes) and washed 2x with PBS-Tx (PBS + 0,1% Triton X-100 w/v) each time spinning the cells down (3000 g, 4°C, 5 minutes). After the final wash, as much supernatant as possible was removed from the pellet, and the tube was snap frozen in liquid nitrogen and stored at -80°C for the ChIP.

### ChIP protocol

For ChIP a number of tubes corresponding to the appropriate number of fixed larval plasmatocytes were thawed on ice. Samples were then resuspended in ice cold PBS-Tx (PBS + 0,1% Triton X-100 w/v) and pooled. If several different ChIPs were performed in one run, they derived from a common cell pool. If biological replicates of a ChIP were performed in one run, these samples derive from separate cell pools. Pooled fixed plasmatocytes were centrifuged (4500 g, 4°C, 10 minutes) and all supernatant was drained. Cells were then resuspended in RIPA buffer (10 mM Tris-HCl, 140 mM NaCl, 1 mM EDTA, 1% (v/v) Triton X-100, 0.1% (w/v) SDS, 0.1% (w/v) sodium deoxycholate, pH 8.0) supplemented with protease inhibitor cocktail (PhosSTOP) and 20 mM sodium butyrate to a final concentration of 3 larval equivalents of plasmatocytes per µl. Plasmatocytes were then dissociated by passing the cell suspension 10 times through a G26 syringe and incubated for 30 minutes on ice. The chromatin suspension was then split into prechilled Bioruptor microtubes at 100 µl per tube and sonicated in a Bioruptor Pico using 30 s on 30 s off cycles at 4°C. Sonication was performed first for 3 cycles and then 2 times for 4 cycles, vortexing tubes inbetween. Sonicated chromatin samples were then spun down to remove unsheared debris (16000 g, 4°C, 10 minutes) and the supernatant was pooled again into a fresh DNA LoBind tube. From this pool IPs were set according to the required plasmatocyte counts holding a small amount back as input. Volumes were adjusted to 100 µl using RIPA and antibodies were added at appropriate concentrations (see materials). The chromatin antibody mixture was incubated on a rotating wheel at 4°C overnight. The next day a 1:1 mixture of protein A and protein G Dynabeads was washed 3 times using a DynaMag-2. Then the beads were resuspended in a small volume and added to the overnight samples corresponding to 5 µl of each protein A and G beads per sample. Bead-antibody-chromatin samples were incubated on a rotating wheel at 4°C for 4 hours. Beads were pulled down using a DynaMag-2, resuspended in fresh RIPA and washed for 10 minutes on a rotating wheel at 4°C. This washing was repeated 4 times with RIPA500 (10 mM Tris-HCl, 500 mM NaCl, 1 mM EDTA, 1% (v/v) Triton X-100, 0.1% (w/v) SDS, 0.1% (w/v) sodium deoxycholate, pH 8.0), once with LiCl buffer (10 mM Tris-HCl, 250 mM LiCl, 1 mM EDTA, 0.5% (v/v) IGEPAL CA-630, 0.5% (w/v) sodium deoxycholate, pH 8.0) and 2 times with TE buffer (10mM Tris-HCl, 1mM EDTA, pH 8.0), for which washing was limited to 5 minutes. Finally, the beads were resuspended in 100 µl RIPA and the same was done for the untreated input sample. 1 µl of RNase (10 mg/ml) was added to each sample and incubated for 1 h at 37°C. 4 µl 10% SDS and 3 µl Proteinase K (20 mg/ml) was added to each sample and the samples were incubated overnight at 65°C shaking at 1200 rpm. The next day magnetic beads were removed using the DynaMag-2 and DNA was isolated from the supernatant using the Zymo ChIP DNA clean and concentrator kit according to the manual. The DNA was finally eluted in 30 µl ultra-pure water and stored at - 80°C.

### ChIP quality control

ChIP samples were checked for concentration of DNA using 5 µl of ChIP sample with the Qubit DNA high sensitivity kit using the Qubit 2.0 fluorometer. Differential enrichment of genomic regions by ChIP was then confirmed by qPCR. 2 µl of each ChIP sample was diluted across reactions for 8 regions with 2 replicates each and qPCR was performed using the Fast SYBR Green master mix with primers targeted against well characterized genomic regions according to the manufacturer’s instructions. All samples passed these quality control tests.

### ChIP library preparation

ChIP-seq libraries were prepared from the final ChIP samples using the NEBNext Ultra II DNA kit and the NEBNext Multiplex Oligos for Illumina sets according to the manual with one minor adjustment: size selection of the library was performed after PCR amplification. For this, after PCR amplification, 27,5 µl of AMpure XP beads were added to each sample and incubated for 5 minutes. Then, the tubes were placed on a DynaMag-2 and the supernatant was moved to fresh tubes, while the beads were discarded. To the supernatant 22,5 µl AMpure XP beads was added, they were incubated for 5 minutes at room temperature, then pulled down using the DynaMag-2 and the supernatant was discarded.

The beads were washed 2 times on the magnetic rack with 80% EtOH and then resuspended in 17 µl ultra-pure water. After incubating for 2 minutes at room temperature, the magnetic beads were pulled down and the supernatant containing the final libraries was moved to fresh tubes. The libraries were then stored at -20°C.

### Library quantification and pooling

Both, RNA-seq and ChIP-seq libraries were processed the same way for quality control and sequencing. First DNA concentration was determined in all libraries using the Qubit DNA high sensitivity kit. 1 µl of the library was then diluted to 1 ng/µl using EB buffer (10mM Tris-HCl, 0.05% Tween-20, pH 8.0) and run on an Agilent bioanalyzer high sensitivity DNA ChIP. If the library was sufficiently concentrated and no adaptors were visible in the bioanalyzer tracks, library concentrations were determined using the KAPA library quantification kit. All libraries were diluted 1:300000 and run in 2 wells of a MicroAmp plate in a QuantStudio3 system according to the KAPA kit instructions. From the Ct values relative molar abundances of each library were calculated as 2^-Ct^ and pooling ratios were determined. Where necessary, libraries were diluted in EB buffer.

### Library sequencing

Libraries were sequenced either at Max Planck Genome Centre (Cologne), at the Max Planck Institute for Molecular Genetics (Berlin) or at Eurofins MWG.

### RNAseq data mapping and quality control

All sequencing data was transferred from the Max Planck Genome Centre (Cologne), the Max Planck Institute for Molecular Genetics (Berlin) or Eurofins MWG as fastq files that were pre-split according to library indexes with a 0 mismatch de-multiplexing. The quality of the sequencing run was determined using FastQC.

The reference genome fasta sequence file of the Berkeley Drosophila Genome Project assembly dm6 and the related gtf genome annotation file for dm6 of ensembl release 91 (dm6.91) were downloaded from ensembl (www.ensmbl.org) (*39*). A reference genome index was generated using dm6.91 using STAR-2.7.0e (*40*) calling

> STAR --runMode genomeGenerate --runThreadN 10 --genomeDir *TargetDir*
>
> --genomeFastaFiles *SourceDir*/Drosophila_melanogaster.BDGP6.dna.toplevel.fa
>
> --sjdbGTFfile *SourceDir*/Drosophila_melanogaster.BDGP6.91.gtf

The index was used to map the fastq files to the genome using

> STAR --genomeDir *GenomeDir* –readFilesIn *ForwardMate*.fastq.gz *ReverseMate*.fastq.gz
>
> --readFilesCommand gunzip -c --outSAMtype BAM SortedByCoordinate
>
> --alignIntronMin 12 --outFileNamePrefix *OutputDir* --runThreadN 6

An index file was generated for the resulting bam file using samtools (*41*)

> samtools index *TargetBam*

And duplicate reads were removed using Picard tools

> java -Xmx32G -jar picard-2.8.2.jar MarkDuplicates INPUT=*StarOutput.bam*
>
> OUTPUT=*NoDuplicateFile.bam*
>
> METRICS_FILE=*SampleID*_MarkDuplicates_metrics.txt
>
> OPTICAL_DUPLICATE_PIXEL_DISTANCE=2500 CREATE_INDEX=true
>
> TMP_DIR=/tmp REMOVE_DUPLICATES=true ASSUME_SORTED=true

BigWig signal track files were generated from the duplicate free bam file using IGVtools (*42*) and UCSCtools (*43*)

> igvtools count --minMapQuality 30 *NoDuplicateFile.bam* stdout dm6 | wigToBigWig -
>
> clip dm6.chrom.sizes *NoDuplicateFile*.bw

Quality control of RNA-seq mapping was further performed using RSeQC (*44*).

All quality control files for FastQC, STAR mapping, Picard tools and RSeQC were aggregated and visualized using MultiQC (*45*) and all data was checked to make sure the library and sequencing was of good quality.

Once a data set passed quality control, the gene level read counts were determined from duplicate free bam files using the subread package (*46*)

> featureCounts -p -s 2 -t exon -g gene_id -a Drosophila_melanogaster.BDGP6.91.gtf -o
>
> *RunID*.count *NoDup1.bam NoDup2.bam NoDup3.bam …*

These read counts were then used for further analysis.

### RNAseq differential expression analysis

The gene level read counts were then loaded into R. For PCA analysis the matrix of gene level read counts was transformed using the DESeq2 (*47*) rlog function, from which the 1000 most variant genes were selected, PCA analysis was performed using the stats package prcomp function and visualized using the plot3D package. For differential expression analysis, gene level read counts were processed using the edgeR package with the quasi-likelihood general linear model approach (*48, 49*) according to the manual.

The resulting data was plotted using ggplot2. For heatmaps DESeq2 (*47*) rlog transformed read counts were transformed to z-scores using the R base package scale function and plotted using the ComplexHeatmaps package.

Enrichment testing of gene sets was performed by a hypergeometric test using the stats package phyper function. Effect size was determined as fold enrichment over the expected gene count under the null hypothesis.

### Single Cell data Analysis

Singe cell data of Tattikota et al. (*19*) was downloaded from www.flyrnai.org/scRNA/blood/ as counts per cell cluster. From these counts, cpm were calculating by dividing by the total number of counts in individual cell clusters. Gene state annotation was carried over from our ChIP-Seq analysis to divide genes by state.

### ChIP data mapping and quality control

As for RNA-seq, all sequencing data was transferred from the Max Planck Genome Centre (Cologne) or at the Max Planck Institute for Molecular Genetics (Berlin) as fastq files that were pre-split according to library indexes with a 0 mismatch de-multiplexing. The quality of the sequencing run was determined using FastQC. A reference genome index was generated as above (see RNA-seq).

The resulting index was used to map the fastq files to the genome using

> STAR --genomeDir *GenomeDir* –readFilesIn *ForwardMate*.fastq.gz *ReverseMate*.fastq.gz
>
> --readFilesCommand gunzip -c --outSAMtype BAM Unsorted –alignIntronMax 1
>
> --outFileNamePrefix *OutputDir* --runThreadN 6

Next, the bam files were sorted and an index file was generated for the resulting bam file using samtools (*41*)

> samtools sort *TargetBam*
>
> samtools index *TargetBam*

For quality control insert size and alignment metrics were collected using Picard tools

> java -Xmx32G -jar picard-2.8.2.jar CollectAlignmentSummaryMetrics I=*StarOutput.bam*
>
> O=*AlignmentMetrics.txt* R=Drosophila_melanogaster.BDGP6.dna.toplevel.fa &
>
> java -Xmx32G -jar picard-2.8.2.jar CollectInsertSizeMetrics I=*StarOutput.bam*
>
> O=*InsertSizeMetrics.txt* P=*InsertSizeHistogram.pdf*

Duplicate reads were removed using Picard tools

> java -Xmx32G -jar picard-2.8.2.jar MarkDuplicates INPUT=*StarOutput.bam*
>
> OUTPUT=*NoDuplicateFile.bam*
>
> METRICS_FILE=*SampleID*_MarkDuplicates_metrics.txt
>
> OPTICAL_DUPLICATE_PIXEL_DISTANCE=2500 CREATE_INDEX=true
>
> TMP_DIR=/tmp REMOVE_DUPLICATES=true ASSUME_SORTED=true

All quality control files for FastQC, STAR mapping and Picard tools were aggregated and visualized using MultiQC (*45*) and all data was checked to make sure the library and sequencing was of good quality.

BigWig signal track files were generated from the duplicate free bam file using IGVtools (*42*) and UCSCtools (*43*)

> igvtools count --minMapQuality 30 *NoDuplicateFile.bam* stdout dm6 | wigToBigWig -
>
> clip dm6.chrom.sizes *NoDuplicateFile*.bw

These BigWig files were used to visualize genome tracks in IGV (*42*). Publication figures were generated using SparK (*50*).

For ChIP heatmaps, PCAs and gene signal profiles, ChIP bam files were processed using a local version of Deeptools (*51*) according to the manual, data was transferred to R and visualized using ggplot2.

For peak calling, bam files were used to call macs2 (*52*) (version 2.2.7.1) using the callpeak command with the --BROAD option. For analysis the .boradPeak files were used, where highly enriched regions were defined by q <= 10^−40^, and all peaks use the default cutoff of q <= 0.05.

### Data extraction for model fitting

For applying the binomial expectation maximization algorithm to gene level ChIP signals, gene level read counts from mapped duplicate free ChIP-seq bam files were counted using the subread package (*46*)

> featureCounts -BCpO -Q 30 -T 4 -t exon -g gene_id -a
>
> Drosophila_melanogaster.BDGP6.91.gtf -o *RunID*.count *NoDup1.bam NoDup2.bam*
>
> *NoDup3.bam …*

To extract bin level read counts from ChIP-seq bam files the bamsignals bamProfile function with a bin size of 200 was done in R.

The resulting counts were processed into 2 data matrixes, 1 for ChIP signal and 1 for input signal, in which each row represents a gene or bin and each column represents a library or sample.

### The ClassFinder algorithm

In the ClassFinder algorithm we implemented a multivariate binomial mixture model which is fitted using expectation maximization. Others (*53, 54*) have proposed, that ChIP-seq count data can be modeled using a binomial mixture. Here individual background and foreground mixture classes follow a binomial distribution:

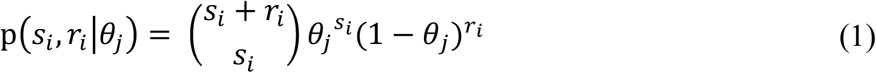

With *i* = 1, …, *N* the number of bins chosen, *s*_*i*_ and *r*_*i*_ the number of reads in the i-th ChIP or input bin respectively, and *θ*_*j*_the probability of “drawing” a ChIP read. In the simple two-component model j can be 1 (background) or 2 (enriched) and *θ*_*j*_ is the same for every bin in the genome which belongs to component j. However, the information whether a genomic region or bin is truly enriched is hidden and figuring out the assignment to the background or enriched class would be the purpose of ChIP-seq peak calling.

Therefore, we must find a way to determine what is a good combination of estimates for both the binomial probabilities θ_j_ and the assignments of the bins. For ease of notation, we will call the true (unknown) assignment of the i-th bin to a component *z*_*i,j*_, which is 1 if the i-th bin is in component j and else 0. Then the likelihood function can be derived from the binomial probability (1)

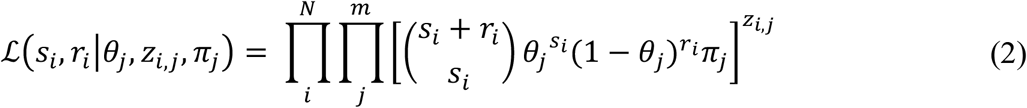

where *π*_*j*_ is the fraction of bins belonging to component *j* and *j* = 1, …, *m* the number of classes used. Now a model can be found that contains the unknown parameters *θ*_*j*_, *z*_*i,j*_ and *π*_*j*_ such that it maximizes the likelihood function (2). This approach is also called maximum likelihood estimation and is useful, because it finds the best possible model, given the mathematical assumptions, that explains the observed data. However, because of the unobserved variables *z*_*i,j*_ the likelihood function cannot be maximized directly.

Therefore, the EM algorithm is used to estimate class assignment. First, the likelihood function is log-transformed (2). For the log-likelihood, we find the simpler equation

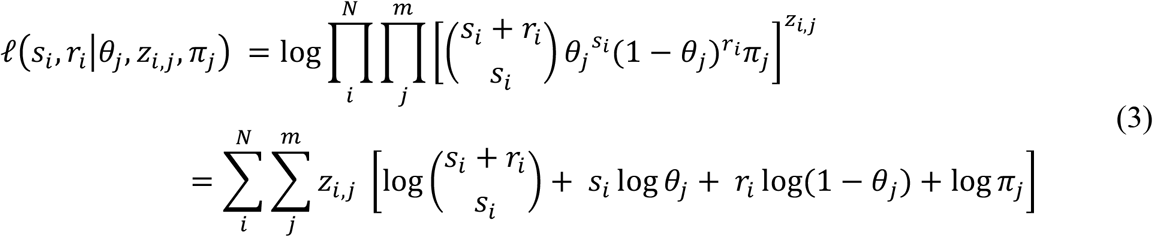

For the expectation-maximization algorithm *z*_*i,j*_ is replaced by the expected value for *z*_*i,j*_ *E*[*z*_*i,j*_|*s*_*i*_, *r*_*i*_, *θ*_*j*_, *π*_*j*_]= *E*_*i,j*_, which results in the expected value of the likelihood function given the current set of parameters:

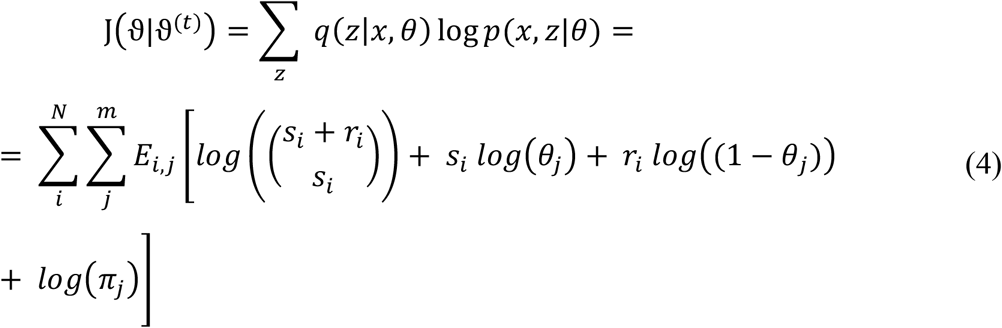

Now the EM algorithm optimizes the likelihood as follows: The model is initialized with a random selection of *θ*_*j*_ and *π*_*j*_ from the space of possible parameters. Then, the function is iteratively optimized by performing

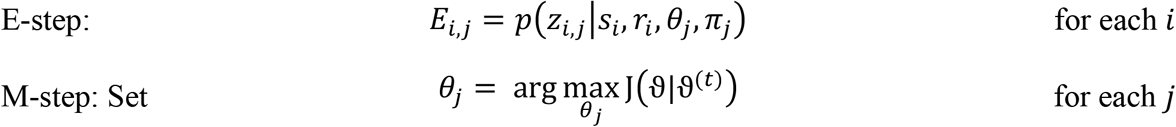

This process is continued iteratively until the change in the expected value of the likelihood function (4) drops below a chosen threshold. As such the EM algorithm can be considered a gradient ascend in which we iteratively maximize J with respect to *E*_*i,j*_ and to the parameters *θ*_*j*_ and *π*_*j*_. Others have demonstrated that such iterative accent at least converges to a local stationary point in J, such a point can however also be a saddle point or, rarely, a local minimum (*55, 56*). Most often, however, for well-behaved likelihood functions this will find at least a local maximum (*56*). In order to maximize the chances of finding a global maximum, the EM-algorithm can impute a set of estimates several times from a different set of randomly chosen starting parameters. When these converged to their local maxima, the results which had the highest likelihood can be chosen ((2) or (3)) as the best model to explain the data.

Therefore, in the t-th iteration of the algorithm, we calculate the E-step with fixed *θ*_*j*_ and *π*_*j*_:

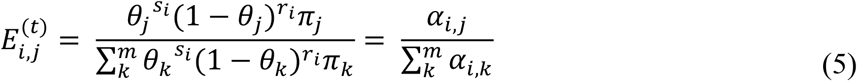

and then maximize the parameter 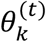 while keeping the *E*_*i,j*_ fixed in the M-step:

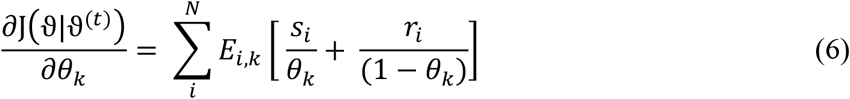

Which solves to

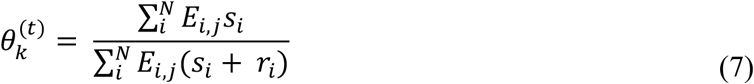

In a similar fashion, we can optimize for *π*_*j*_:

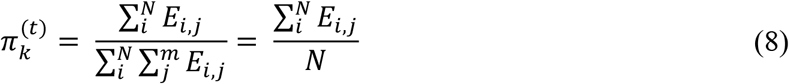

This models the single dimensional case as described elsewhere (*53*). However, we wanted to combine information from multiple replicates. Therefore, we generalized the above mixture model for cases of multivariate binomial mixtures with shared class assignment. For *d* ChIP input pairs (*d* dimensions) the i-th genomic bin the counts *s*_*i*_ and *r*_*i*_ can be transformed into vectors ***s***_*i*_ and ***r***_*i*_ such that ***s***_*i*_, ***r***_*i*_ ∈ ℕ^*d*^ and the probabilities *θ*_*j*_ can be transformed into vectors ***θ***_*j*_ ∈ ℝ^*d*^. We can then extend (1) to:

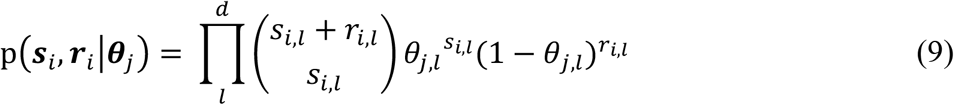

with *l* = 1, …, *d* being the individual dimensions or pairs of samples. Such a probability function is still well normalized since for any vector **c** ∈ ℕ^*d*^ it is true that for the sum over all tuples of ***s, r*** such that ***s*** + ***r*** = ***c***

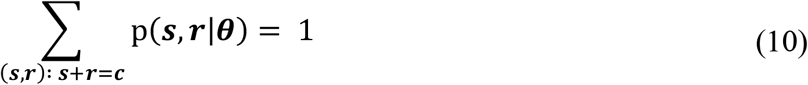

Therefore, the earlier likelihood function (2) can be modified by accounting for the multi-dimensional case:

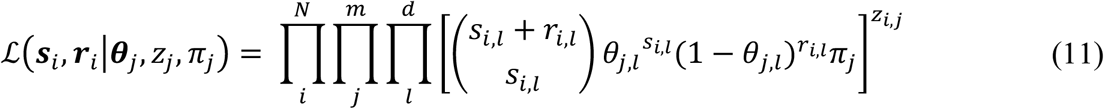

One could apply the brute force approach of finding solutions to every parameter by adding the additional index *l*, but this makes computation inefficient for higher-dimensional cases. By using the representations:

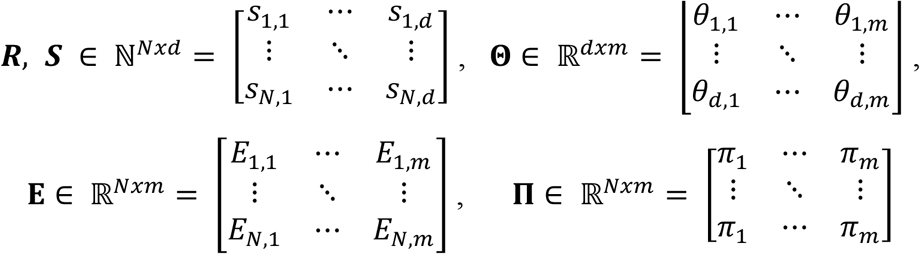

we can find the following simplifications. For the E-step we use

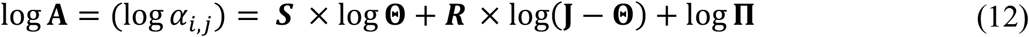

with **J** being a matrix of ones. From there we adapt from (5):

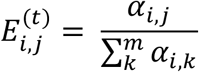

For the M-step, the calculation of *π*_*j*_ is described in (8) and requires no modifications, since the number of dimensions *d* does not apply to this parameter. For the binomial probabilities *θ*_*j,l*_ we calculate

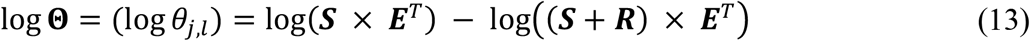

Using formulas (12) and (5) for the E-step and formulas (8) and (13) for the M-step, the EM-algorithm can then be applied to find the estimators for the binomial probabilities *θ*_*j,l*_ as well as the probabilities *E*_*i,j*_ with which each genomic region *i* belongs to a component *j*. This multivariate binomial mixture model is implemented in the ClassFinder algorithm. The model is supplied with the total number of reads in each bin ***N*** = ***R*** + ***S*** ∈ ℕ^*Nxd*^, the number of reads in the ChIP sample ***S*** ∈ ℕ^*Nxd*^ and the number of classes *m*. Further, the number of attempted imputations can be specified (here we used 10). We found that most starting parameters will arrive at the same MLEs, but on highly complex samples that could contain a larger number of local maxima, it might be useful to increase this further. The multithreading, in this case, was simply implemented by running each imputation as a separate thread.

Each imputation starts by randomly sampling **Θ** = *θ*_*j,l*_ *ϵ* [0,1] and *π*_*j*_ *ϵ* [0,1]. Then, iteratively, model estimates are found: First, in the E-step, the expectations are calculated according to formulas (12) and (5), then in the M-step the maximum likelihood estimator of the parameters *θ*_*j,l*_ and *π*_*j*_ are calculated according to (8) and (13). Now the improvement in the log likelihood (see (11)) needs to be calculated to determine if the model has converged. Because the log-likelihood is essentially

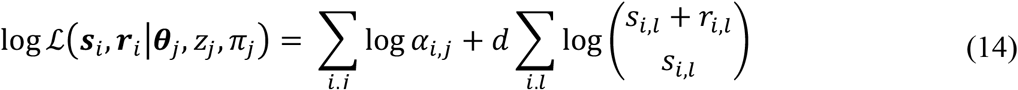

the difference in log-likelihood between iterations t and t+1 are easily found by

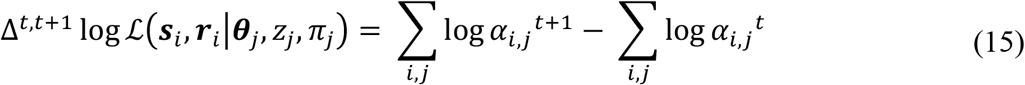

which can be derived from (12). Once that difference is smaller than the set threshold the model is considered converged and saved.

After all imputations are done all resulting models are compared for their log-likelihood and the model with the largest likelihood is chosen as the best model to explain the data. Last, each of the bins on which ChIP and input reads were counted is assigned to one of the components *j* by choosing that component with the highest expectation in the final best model (maximum posterior expectation).

This algorithm was used to fit the read count outputs of the featureCounts or bamsignal functions.

For assignment of 200-bp windowed chromatin states to genes, overlaps with exons were identified using the GenomicRanges (*57*) findOverlap function. States were assigned to genes by largest overlap.

### Class Finder on External data Sets

For analysis of external datasets, we identified H3K27me3 datasets that used individual *Drosophila melanogaster* tissues or cell lines (as opposed to whole animals) on the European Nucleotide Archive. The datasets used were:

| Accession | Cell types | Author | link |
| --- | --- | --- | --- |
| PRJNA120421 | cell line | Gan et al., 2010 | https://www.ncbi.nlm.nih.gov/pmc/articles/PMC2884545/?tool=EBI |
| PRJNA157885 | neuronal cells | Henry et al., 2012 | https://www.ncbi.nlm.nih.gov/pmc/articles/PMC3479168/?tool=EBI |
| PRJNA159935 | cell line | Van Bortle et al., 2012 | https://www.ncbi.nlm.nih.gov/pmc/articles/PMC3483547/?tool=EBI |
| PRJNA176964 | cell line | Herz et al., 2021 | https://www.ncbi.nlm.nih.gov/pmc/articles/PMC3521626/?tool=EBI |
| PRJNA239010 | parasegments | Bowman et al., 2014 | https://www.ncbi.nlm.nih.gov/pmc/articles/PMC4139060/?tool=EBI |
| PRJNA298922 | imaginal discs | Loubiere et al., 2016 | https://europepmc.org/article/PMC/5407438 |
| PRJNA380775 | testes | Laktionov et al., 2018 | https://epigeneticsandchromatin.biomedcentral.com/articles/10.1186/s13072-018-0183-3 |
| PRJNA534289 | cell line | Heurateau et al., 2020 | https://genomebiology.biomedcentral.com/articles/10.1186/s13059-020-02106-z |
| PRJNA554678 | cell line | Hao et al., 2020 | https://genome.cshlp.org/content/30/11/1570.full |
| PRJNA555353 | neuronal cells | Abdusselamoglu et al., 2019 | https://pubmed.ncbi.nlm.nih.gov/31748204/ |

Raw fastq H3K27me3 and Input data from these studies were downloaded. For paired end samples, mapping was performed as above, for single end samples, deduplication by Picard Tools was omitted. Gene level counts were generated as above. Datasets were further split by tissue where multiple tissues were used in individual studies. This data was used for ClassFinder analysis as above.

### Differential ChIP-seq

For differential ChIP-seq data was mapped as described. Gene level read counts from mapped duplicate free ChIP-seq bam files were counted using the subread package (*46*) as before.

Quantile normalization was performed on either log2 ratio bigwig signals or gene level log2 ratio of counts. Figures were generated using SparK (https://www.biorxiv.org/content/10.1101/845529v1.full)

For plotting of gene level differential modification, quantile normalized log2 ratios of gene read counts were centered to the gene mean and averaged across replicates. From this data Pc-M genes were visualized. Significance was tested using a Mann-Whitney test.

## Code Availability

All code that was used for analysis in this paper is available at GitHub (https://github.com/robertstreeck/PolycombPaperR).

## Data Availability

Raw sequencing data and processed data is available on ArrayExpress:

| <u>Accession</u> | <u>Title</u> |
| --- | --- |
| E-MTAB-10759 | RNA-seq of resting plasmatocytes from *Drosophila melanogaster* third instar larvae |
| E-MTAB-10762 | RNA-seq of *Drosophila* plasmatocyte mosaics which lack PRC2/H3K27me3 |
| E-MTAB-10780 | RNA-seq of plasmatocytes after septic injury with *trxG*-RNAi |
| E-MTAB-10760 | ChIP-seq of *Drosophila melanogaster* OregonR plasmatocytes at 6h post septic injury |
| E-MTAB-10761 | RNA-seq of plasmatocytes after septic injury |
| E-MTAB-10764 | RNA-seq of Hml-SwitchGal4 induced pathway activation in *Drosophila* plasmatocytes |
| E-MTAB-10707 | ChIP-seq of *Drosophila melanogaster* plasmatocytes against various histone modifications |

## Antibodies

| <u>Antigen</u> | <u>Species</u> | <u>use</u> | <u>Per ChIP</u> | <u>Cat. No</u> | <u>Lot</u> | <u>Reference</u> |
| --- | --- | --- | --- | --- | --- | --- |
| H3K27me3 | rabbit polyclonal | ChIP | 0,25 µg | C15410195 | A0821D | Diagenode SA, Belgium |
| CTD4H8 RNA polymerase II | mouse monoclonal | ChIP | 1 µg | 05-623 | 2885186 | Merck Millipore, USA |
| H3K9ac | rabbit polyclonal | ChIP | 0,3 µg | C15410004 | A1435-0012D | Diagenode SA, Belgium |
| H3K9me3 | rabbit polyclonal | ChIP | 0,2 µg | C15410193 | A1671-001P | Diagenode SA, Belgium |
| H3K4me1 | rabbit polyclonal | ChIP | 0,15 µg | C15410194 | A1862D | Diagenode SA, Belgium |
| H3K27ac | rabbit polyclonal | ChIP | 0,15 µg | C15410196 | A1723-0041D | Diagenode SA, Belgium |
| H3K36me3 | rabbit polyclonal | ChIP | 0,15 µg | C15410192 | A1845P | Diagenode SA, Belgium |
| H3K4me3 | rabbit polyclonal | ChIP | 0,1 µg | ab8580 | GR240214-2 | Abcam, UK |
| H3K27me2/me3 | mouse monoclonal | ChIP | 0,15 µg | 39535 | 33311011 | Active Motif, Belgium |

## Fly stocks

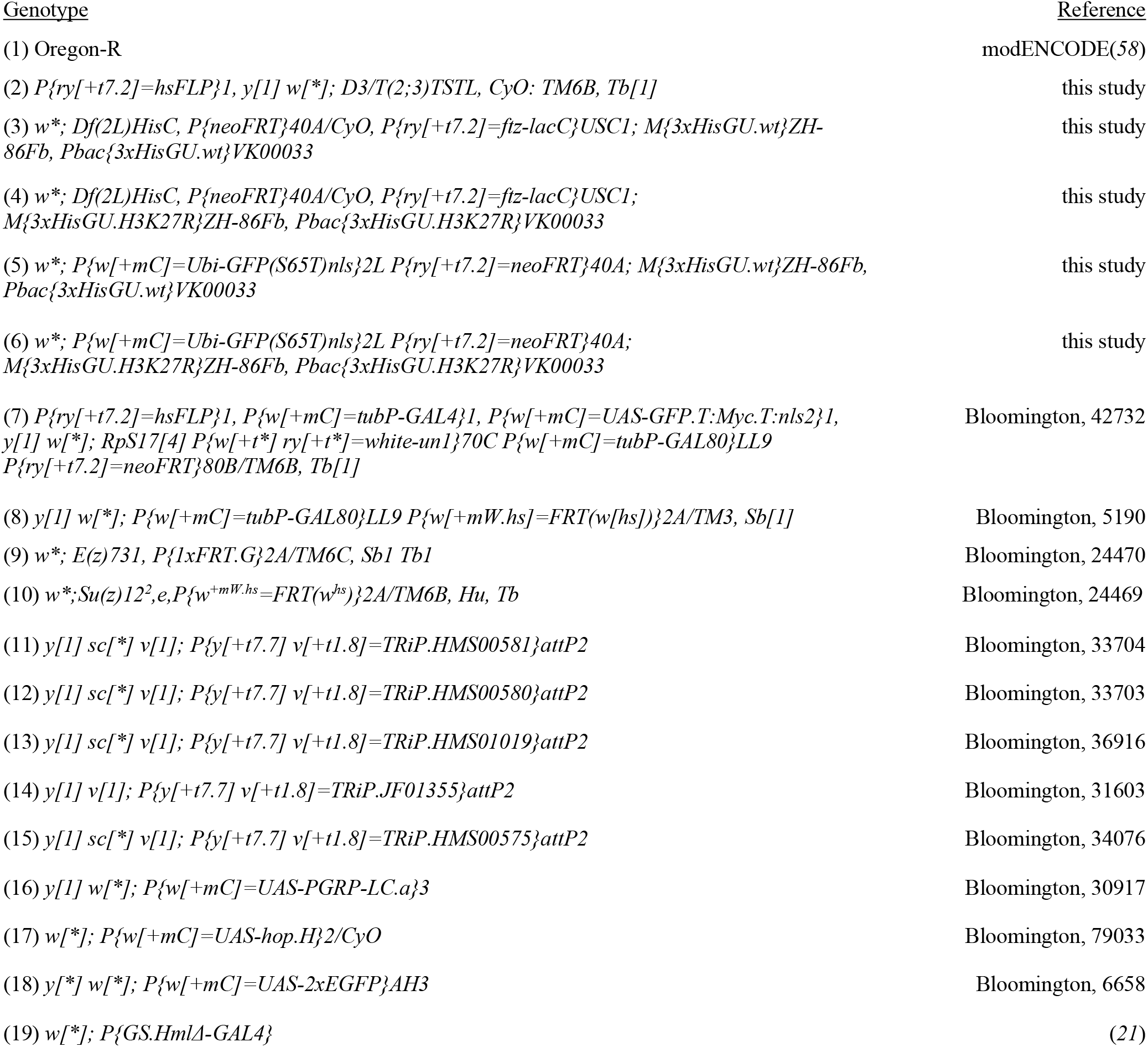

## Acknowledgments

We thank H. Jaspers for fly strains and A. Zychlinsky for comments on the manuscript.

## Author contributions

Conceptualization: RS, AH

Methodology: RS, HNS

Investigation: RS, HNS, AH

Visualization: RS, AH

Supervision: AH

Writing – original draft: RS, AH

Writing – review & editing: RS, HNS, AH

## Data and materials availability

All data are available in the main text or the supplementary materials.

## Supplementary Materials

Figs. S1 to S17

