## Supplementary Figures S1-S17 for "Reversible repression of inducible genes by Polycomb Repressive Complex 2 and H3K27me3 in Drosophila melanogaster"

### Figure S1

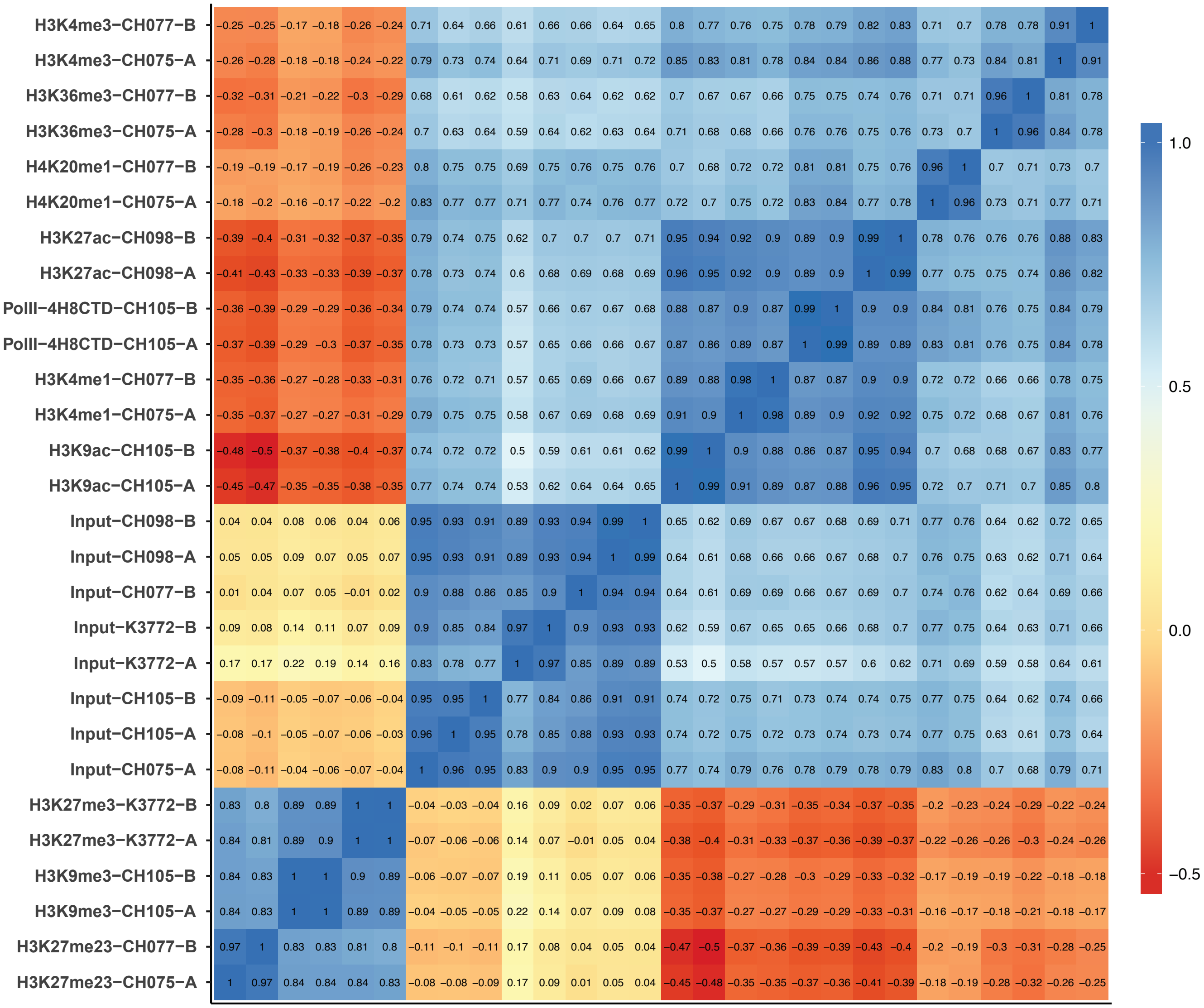

H3K4me3−CH077−B
H3K4me3−CH075−A
H3K36me3−CH077−B
H3K36me3−CH075−A
H4K20me1−CH077−B
H4K20me1−CH075−A
H3K27ac−CH098−B
H3K27ac−CH098−A
PolII−4H8CTD−CH105−B
PolII−4H8CTD−CH105−A
H3K4me1−CH077−B
H3K4me1−CH075−A
H3K9ac−CH105−B
H3K9ac−CH105−A
Input−CH098−B
Input−CH098−A
Input−CH077−B
Input−K3772−B
Input−K3772−A
Input−CH105−B
Input−CH105−A
Input−CH075−A
H3K27me3−K3772−B
H3K27me3−K3772−A
H3K9me3−CH105−B
H3K9me3−CH105−A
H3K27me23−CH077−B
H3K27me23−CH075−A
1.0
0.5
0.0
−0.5

**Figure S1. Plasmatocyte ChIP-seq replicate correlation demonstrates high reproducibility.**

Heatmap of Spearman correlation of ChIP-seq samples. ChIP-seq was performed in duplicates from primary, resting *Drosophila* plasmatocytes. From the resulting sequencing data, correlation analysis was performed using Deeptools. Mapped reads from each individual ChIP-seq replicate were binned using 10kb windows and pairwise Spearman correlation coefficients calculated. Experiment indices (CH*** or K****) indicate experimental batches and final letters (A or B) indicate replicates; color scale indicates correlation coefficients.

### Figure S2

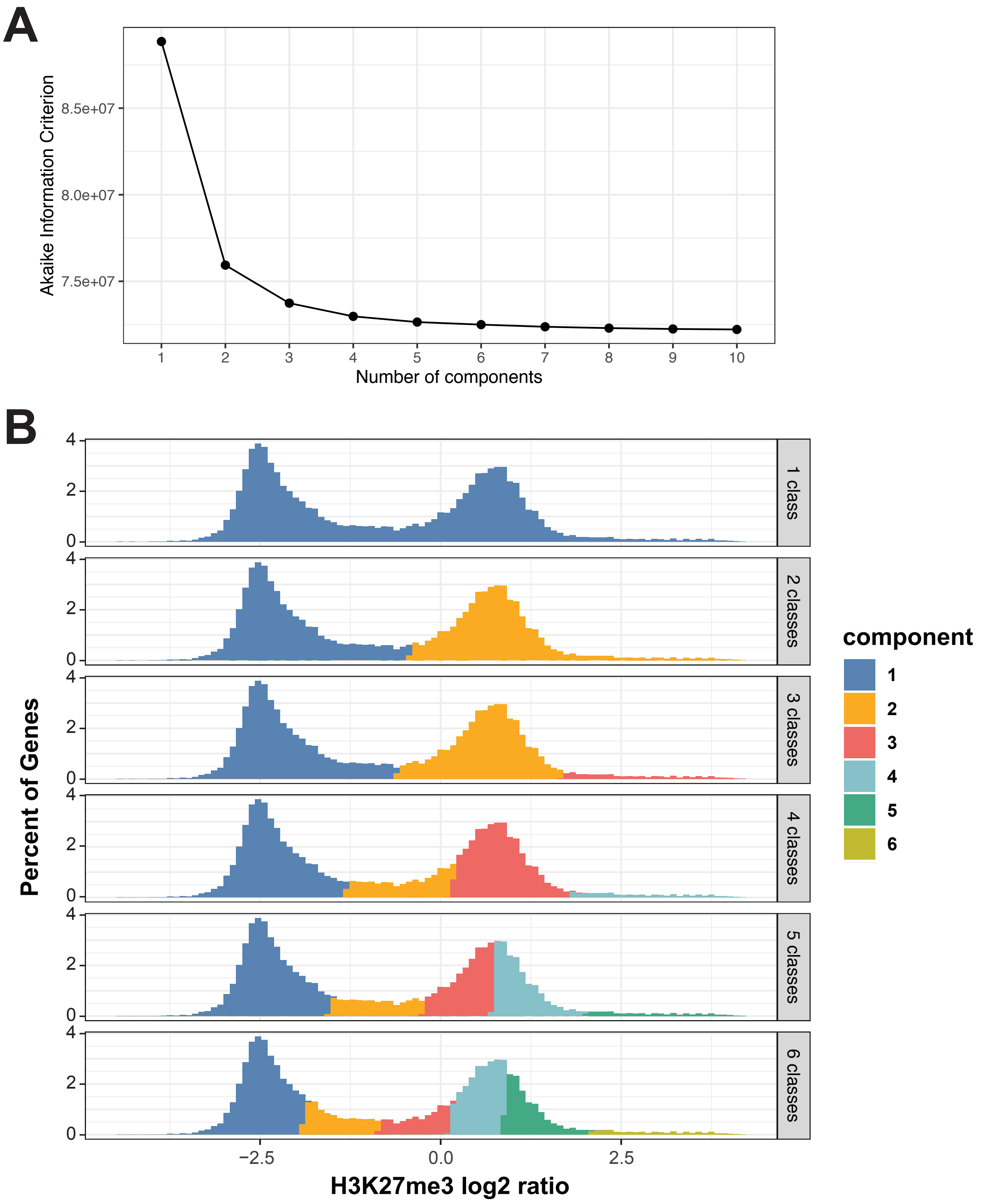

A
Akaike Information Criterion
8.5e+07
8.0e+07
7.5e+07
1
2
3
4
5
6
7
8
9
10
Number of components
B
Percent of Genes
4
2
0
1 class
2 classes
3 classes
4 classes
5 classes
6 classes
−2.5
0.0
2.5
H3K27me3 log2 ratio
component
1
2
3
4
5
6

**Figure S2. Gene-level H3K27me3 is well modeled by a 3-class mixture.**

(**A**) Akaike Information Criterion (AIC) of ClassFinder fits to gene-level H3K27me3 with different numbers of classes (components). H3K27me3 signal was quantified by counting ChIP-seq and Input reads over gene exons using featureCounts. Counts were clustered into 1-10 classes by multivariate binomial mixture modelling using ClassFinder (see Material and Methods). AIC of these models drops quickly up to 3 classes. (**B**) Histograms of all genes ordered by gene-level H3K27me3 signal as log2 ratio over Input (see Fig. 1B). Gene assignment of models fit in (A) for 1-6 classes. The 3-class model well describes the strong bimodal distribution while accounting for the long tail at high H3K27me3 signal. The 4-class model describes the genes intermediate between non-Pc and Pc-M as an additional class, and a 5-class model splits the Pc-M peak in a way that appears unreasonable given the distribution.

### Figure S3

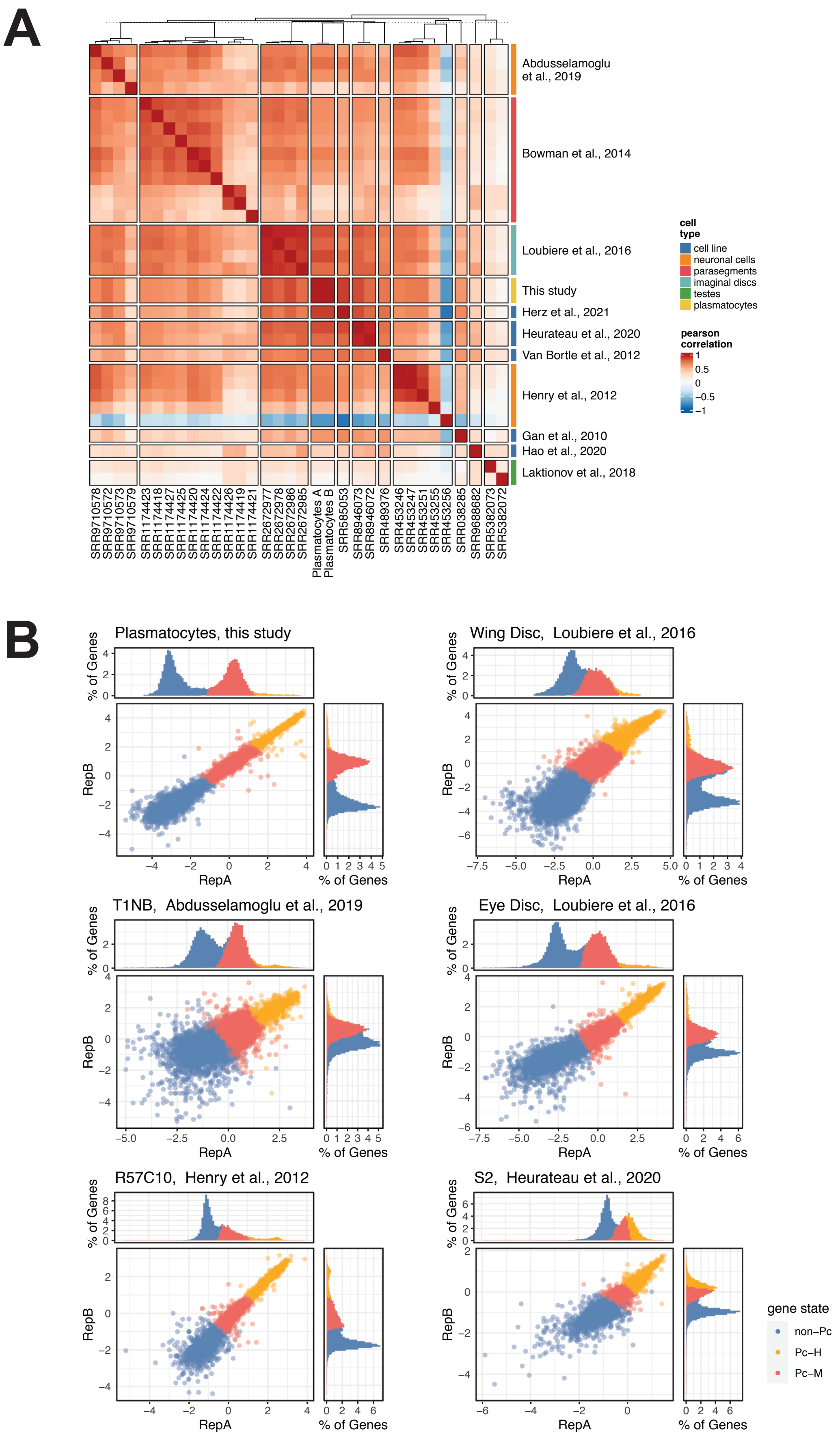
A
Abdusselamoglu et al., 2019
Bowman et al., 2014
Loubiere et al., 2016
This study
Herz et al., 2021
Heurateau et al., 2020
Van Bortle et al., 2012
Henry et al., 2012
Gan et al., 2010
Hao et al., 2020
Laktionov et al., 2018
cell type
cell line
neuronal cells
parasegments
imaginal discs
testes
plasmatocytes
pearson correlation
Plasmatocytes A
Plasmatocytes B
B
Plasmatocytes, this study
Wing Disc, Loubiere et al., 2016
T1NB, Abdusselamoglu et al., 2019
Eye Disc, Loubiere et al., 2016
R57C10, Henry et al., 2012
S2, Heurateau et al., 2020
% of Genes
RepA
RepB
gene state
non–Pc
Pc–H
Pc–M

**Figure S3. Comparable H3K27me3 ClassFinder models can be derived for other *Drosophila* datasets.**

(**A**) Gene-level signal correlation in H3K27me3 datasets. Raw data of H3K27me3 ChIP-seq datasets (see table in Material and Methods) was retrieved from the European Nucleotide Archive, mapped and counted as described in Material and Methods. Pairwise Pearson correlation coefficients were calculated and plotted as a heatmap. Heatmap colors indicate correlation coefficients, colors on the right indicate sample origin with the study reference beside it, and bottom IDs identify the sample by ENA reference. (**B**) ClassFinder models of external H3K27me3 data. From all samples in (A), we selected those with Spearman's correlation coefficient $r > 0.7$ between biological replicates and generated a 3-class model using ClassFinder. Inner scatter plots show gene-level H3K27me3 log2 ratio over Input for each replicate, colored by gene class, while the top and right plots show the marginal distributions of the signals. The data from primary cells of imaginal discs (wing discs and eye discs) as well as neuronal tissues (type 1 neuroblast (T1NB) and R57C10 positive cells) show a comparable distribution to our plasmatocyte sample (top left). S2 cells exhibit a distinct bimodal distribution that lacks the Pc-H tail.

### Figure S4

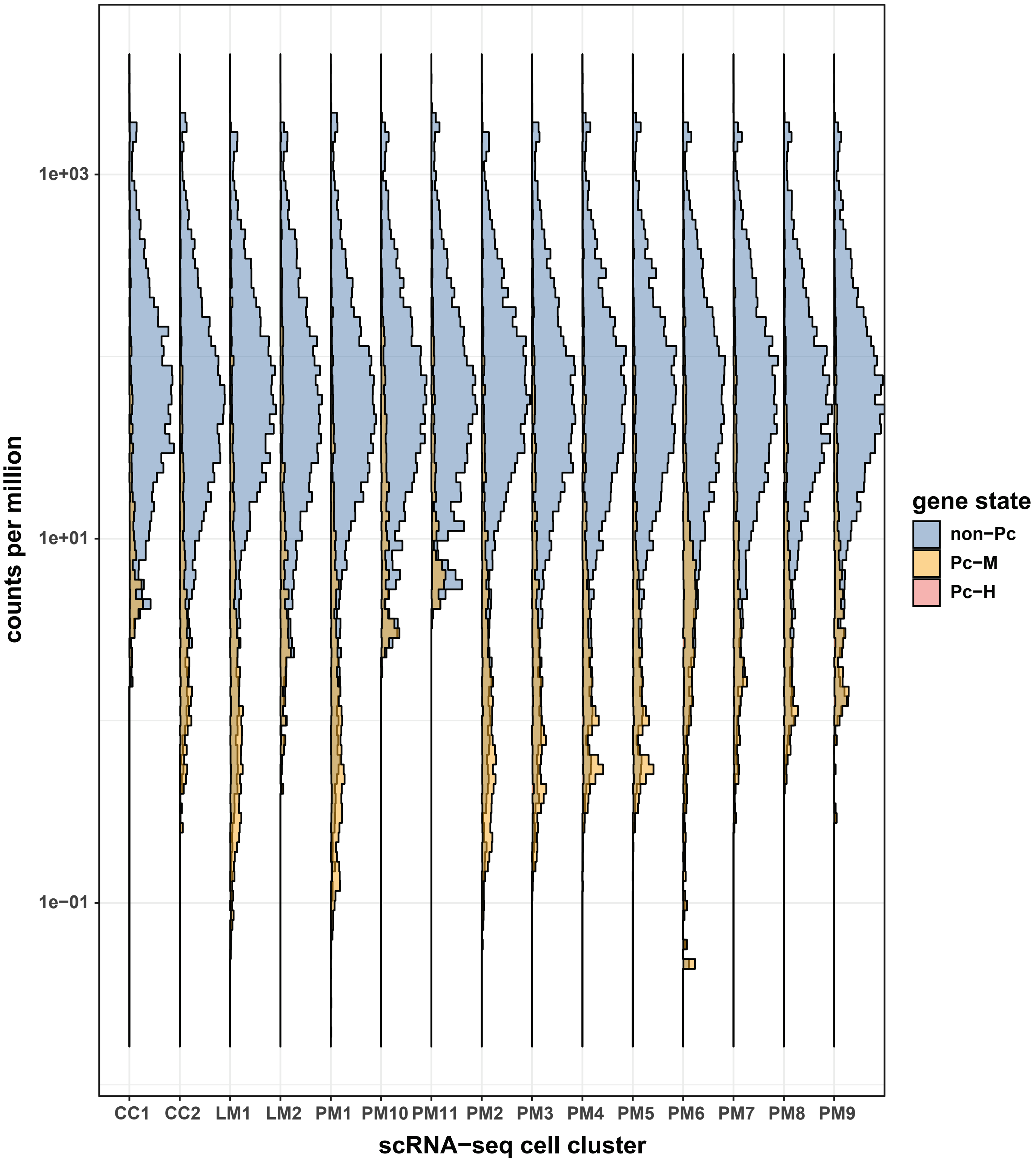

counts per million
1e+03
1e+01
1e-01
CC1
CC2
LM1
LM2
PM1
PM10
PM11
PM2
PM3
PM4
PM5
PM6
PM7
PM8
PM9
scRNA−seq cell cluster
gene state
non−Pc
Pc−M
Pc−H

**Figure S4. Pc-M genes are weakly expressed or not detected across single cell hemocyte clusters.**

Singe cell data of Tattikota *et al.* was retrieved from www.flyrnai.org/scRNA/blood/ as counts per cell cluster and transformed into counts per million. The relative expression strength in counts per million across each scRNA-seq cluster was then plotted with genes grouped by their gene state, as identified by ClassFinder. Pc-M genes are not strongly expressed in any individual cell cluster, indicating that cell heterogeneity is not the reason for weak expression of Pc-M genes detected in the complete plasmatocyte population.

### Figure S5

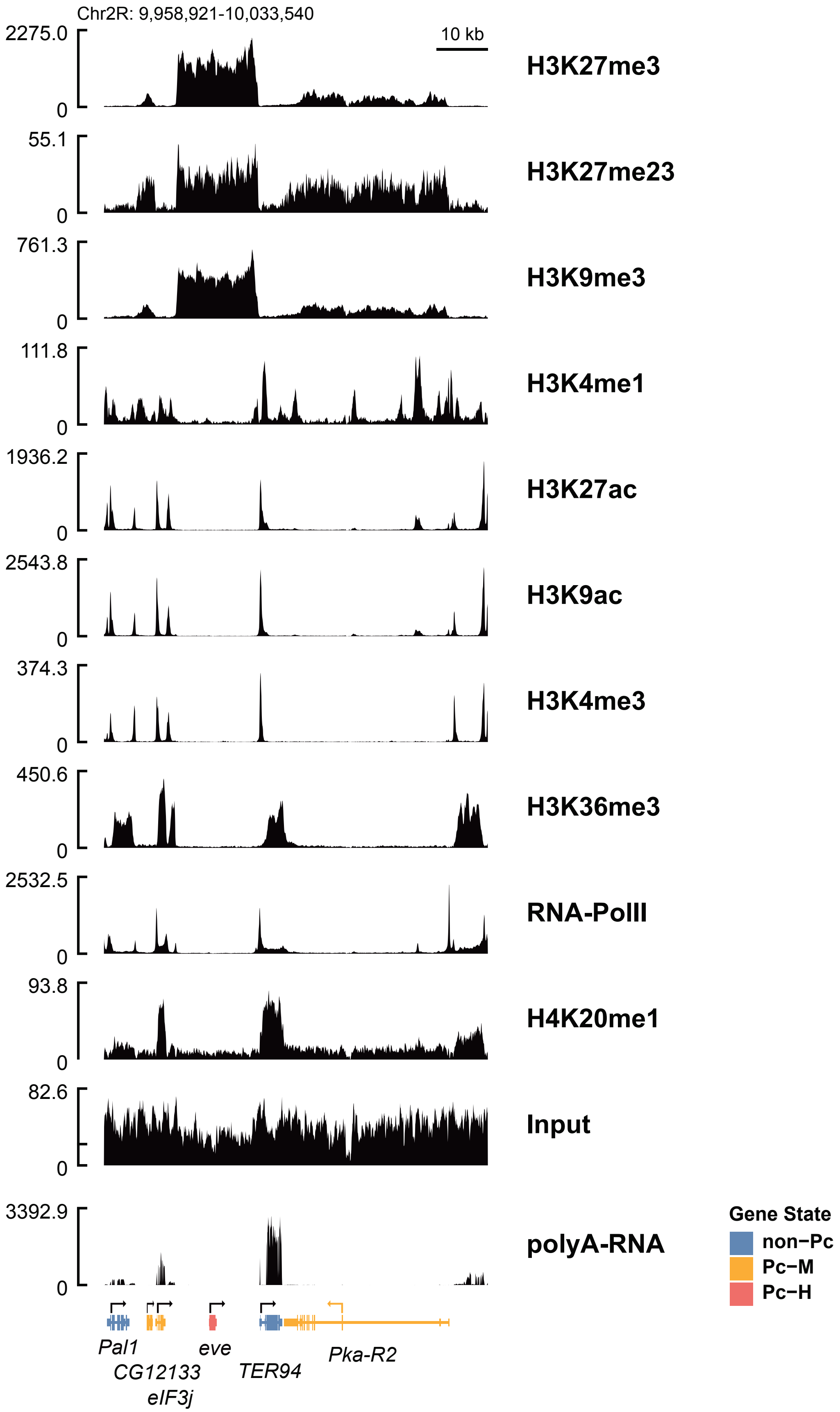

Chr2R: 9,958,921-10,033,540
10 kb
2275.0
0
H3K27me3
55.1
0
H3K27me23
761.3
0
H3K9me3
111.8
0
H3K4me1
1936.2
0
H3K27ac
2543.8
0
H3K9ac
374.3
0
H3K4me3
450.6
0
H3K36me3
2532.5
0
RNA-PolII
93.8
0
H4K20me1
82.6
0
Input
3392.9
0
polyA-RNA
Gene State
non−Pc
Pc−M
Pc−H
Pal1
CG12133
eIF3j
eve
TER94
Pka-R2

**Figure S5. Plasmatocyte ChIP-seq signal around the eve locus across all IP targets.** Raw counts of ChIP-seq (with merged replicates) for the region of Chr2R: 9,958,921-10,033,540. Right labels indicate ChIP targets (see Material and Methods for protocol and antibody details), left shows read counts. Genes are marked at the bottom, colored by their respective gene state. While non-Pc genes show the previously described distribution of marks associated with active transcription, Pc-M and Pc-H lack these modifications. Instead, they carry H3K27me3 and other repressive marks, but at distinct levels.

### Figure S6

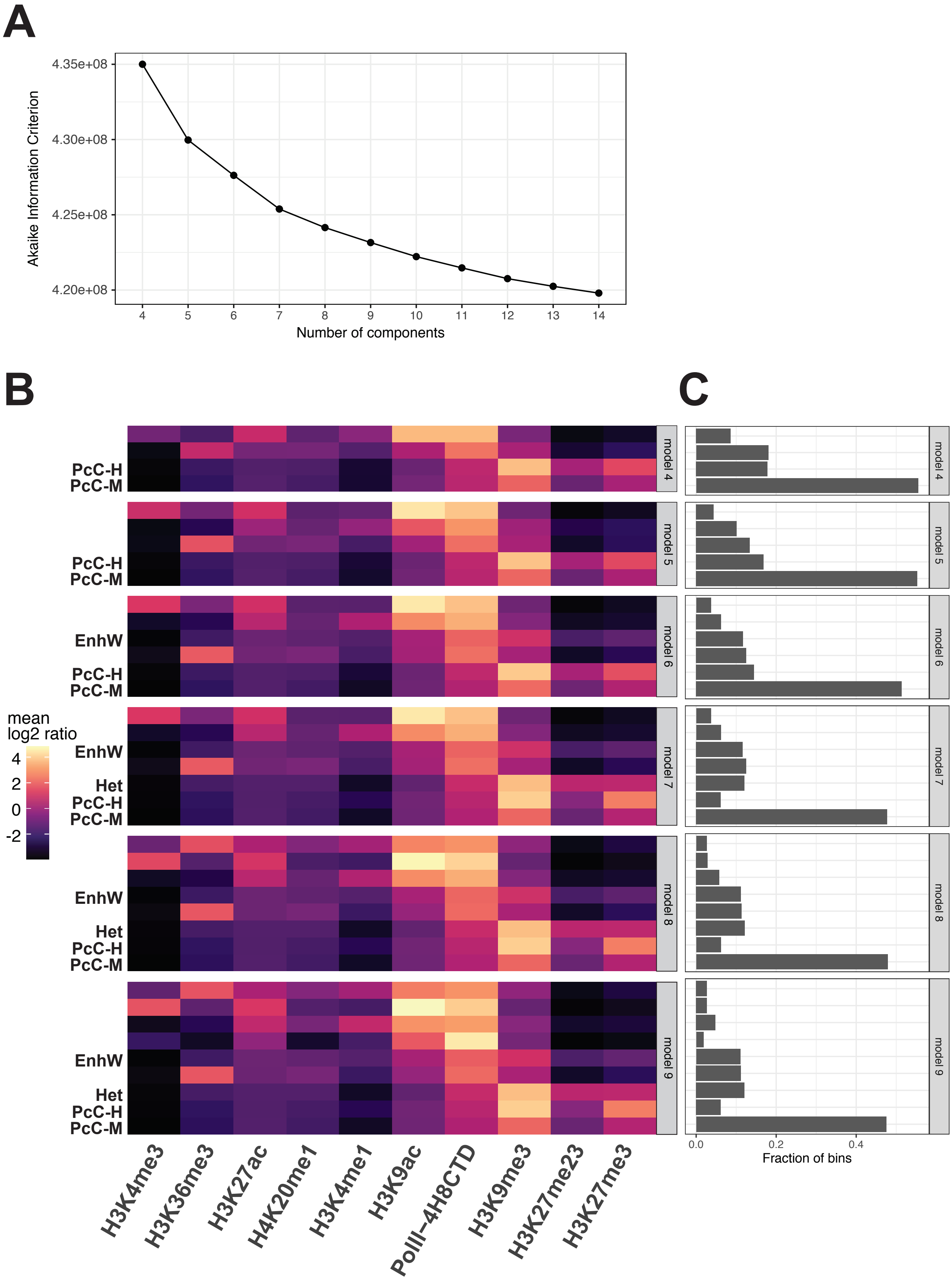

A
4.35e+08
4.30e+08
4.25e+08
4.20e+08
Akaike Information Criterion
Number of components
B
C
PcC-H
PcC-M
EnhW
Het
model 4
model 5
model 6
model 7
model 8
model 9
mean
log2 ratio
4
2
0
-2
H3K4me3
H3K36me3
H3K27ac
H4K20me1
H3K4me1
H3K9ac
PolII–4H8CTD
H3K9me3
H3K27me23
H3K27me3
0.0
0.2
0.4
Fraction of bins

**Figure S6. Chromatin states based on assayed marks are well described by a 7-state ClassFinder model.**

(**A**) Akaike Information Criterion (AIC) at different number of classes (components) in the ClassFinder model. ChIP-seq signal for all assayed marks was quantified for 200 base-pair bins and used to fit ClassFinder models at different numbers of mixture components. AIC decreases across all tested mixtures, with an inflection in decrease at 7 classes. (**B**) Signal heatmaps for 4-9 class models. From the extracted models, mean log2 ratio signals for each assayed mark (below heatmap) were derived from bin level signal. Across all models, Polycomb chromatin is separated into PcC-M and PcC-H states. At 7 classes, a distinct heterochromatic state (Het) appears. Increasing the number of classes beyond 7 further differentiates states associated with active marks. (**C**) Number of bins in each class shown in (B). The PcC-M state covers most of the genome at 4 classes and decreases slightly in prevalence at 6 components (separation of weak enhancer state) and 7 components (separation of heterochromatic state), but is constant afterwards.

### Figure S7

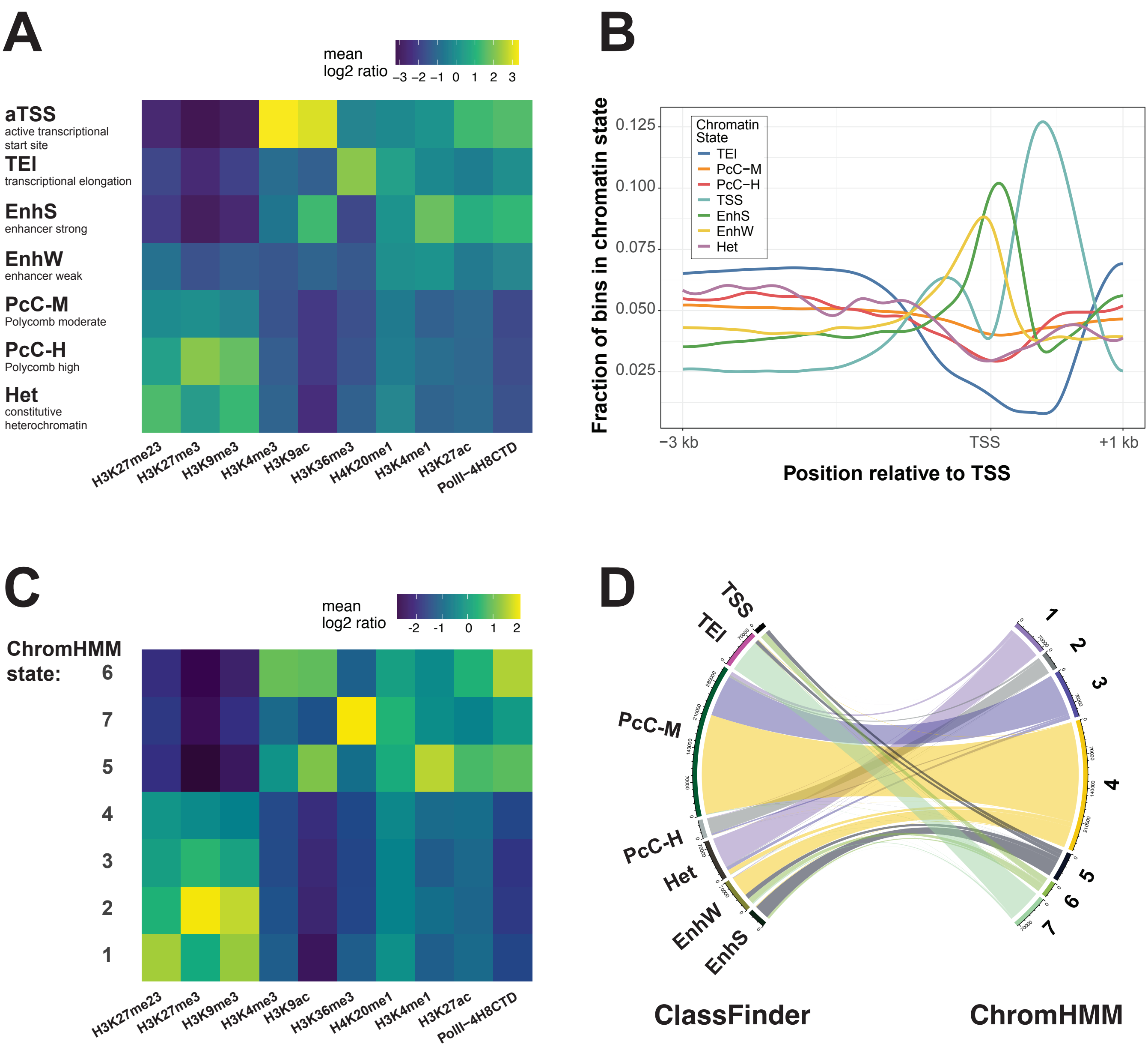

A
mean log2 ratio
−3 −2 −1 0 1 2 3
aTSS
active transcriptional start site
TEI
transcriptional elongation
EnhS
enhancer strong
EnhW
enhancer weak
PcC-M
Polycomb moderate
PcC-H
Polycomb high
Het
constitutive heterochromatin
H3K27me23
H3K27me3
H3K9me3
H3K4me3
H3K9ac
H3K36me3
H4K20me1
H3K4me1
H3K27ac
PolII−4H8CTD
B
Fraction of bins in chromatin state
0.125
0.100
0.075
0.050
0.025
Chromatin State
TEI
PcC−M
PcC−H
TSS
EnhS
EnhW
Het
−3 kb
TSS
+1 kb
Position relative to TSS
C
mean log2 ratio
-2 -1 0 1 2
ChromHMM state:
6
7
5
4
3
2
1
H3K27me23
H3K27me3
H3K9me3
H3K4me3
H3K9ac
H3K36me3
H4K20me1
H3K4me1
H3K27ac
PolII−4H8CTD
D
TSS
TEI
PcC-M
PcC-H
Het
EnhW
EnhS
1
2
3
4
5
6
7
ClassFinder
ChromHMM

**Figure S7. ClassFinder and ChromHMM produce comparable models with some distinctions.**

(**A**) Heatmap of features abundance across a 7-state chromatin model generated by ClassFinder. The 7-state model was fit to all assayed marks (below heatmap) over 200bp windows and the IP target signal was quantified as mean log2 ratio. Assignments of chromatin states are indicated (left of heatmap). This panel is identical to Fig. 1D and shown here to facilitate comparison with supplementary data.

(**B**) Genome coverage of ClassFinder states around transcriptional start sites analyzed as fraction of 200bp genome bins assigned to individual states.

(**A, B**) The active transcriptional start site state (aTSS) is characterized by high level H3K4me3 and H3K9ac, which are both marks found at active promotors. The depletion of this state at the transcriptional start site and its abundance at the +1 nucleosome reflects a typical H3K4me3 pattern. The transcriptional elongation state (TEl) is characterized by high level H3K36me3, which increases distally to the promotor over the gene body of actively transcribed genes. We also identified a 'strong' enhancer state (EnhS) characterized by H3K4me1 and H3K9ac. In addition, we found a chromatin state (EnhW) that shows similar distribution relative to transcriptional start sites but lower abundance of H3K4me1 and H3K9ac, which we therefore assigned as 'weak' enhancer state. The two Polycomb chromatin states (PcC-M, PcC-H) differ primarily in the abundance of H3K27me3 and H3K9me3, with H3K27me3 being more abundant than H3K9me3. In the constitutive heterochromatin state (Het) this ratio is inversed and H3K9me3 is the dominant mark.

(**C**) ChIP-seq signal across 7 states as predicted by ChromHMM (ChromHMM state). ChIP-seq mappings were used to predict chromatin states using ChromHMM (see Material and Methods). From the resulting assignments, ChIP signal was quantified as mean log2 ratio across the class bins and visualized as a heatmap. Similar to the ClassFinder algorithm, distinct active states can be identified for active transcriptional start site (6), transcriptional elongation (7) and enhancers (5). In addition, there are distributions similar to PcC-H (2) and constitutive heterochromatin (1) states, while the remaining two states (3 and 4) are both similar to PcC-M and EnhW states (**D**) Mapping of ChromHMM to ClassFinder 7 state models. Direct comparison of classifications show that the main difference between both models is in the allocation of genome bins into ClassFinder PcC-M/EnhW states versus state 3 and 4 of the ChromHMM model.

### Figure S8

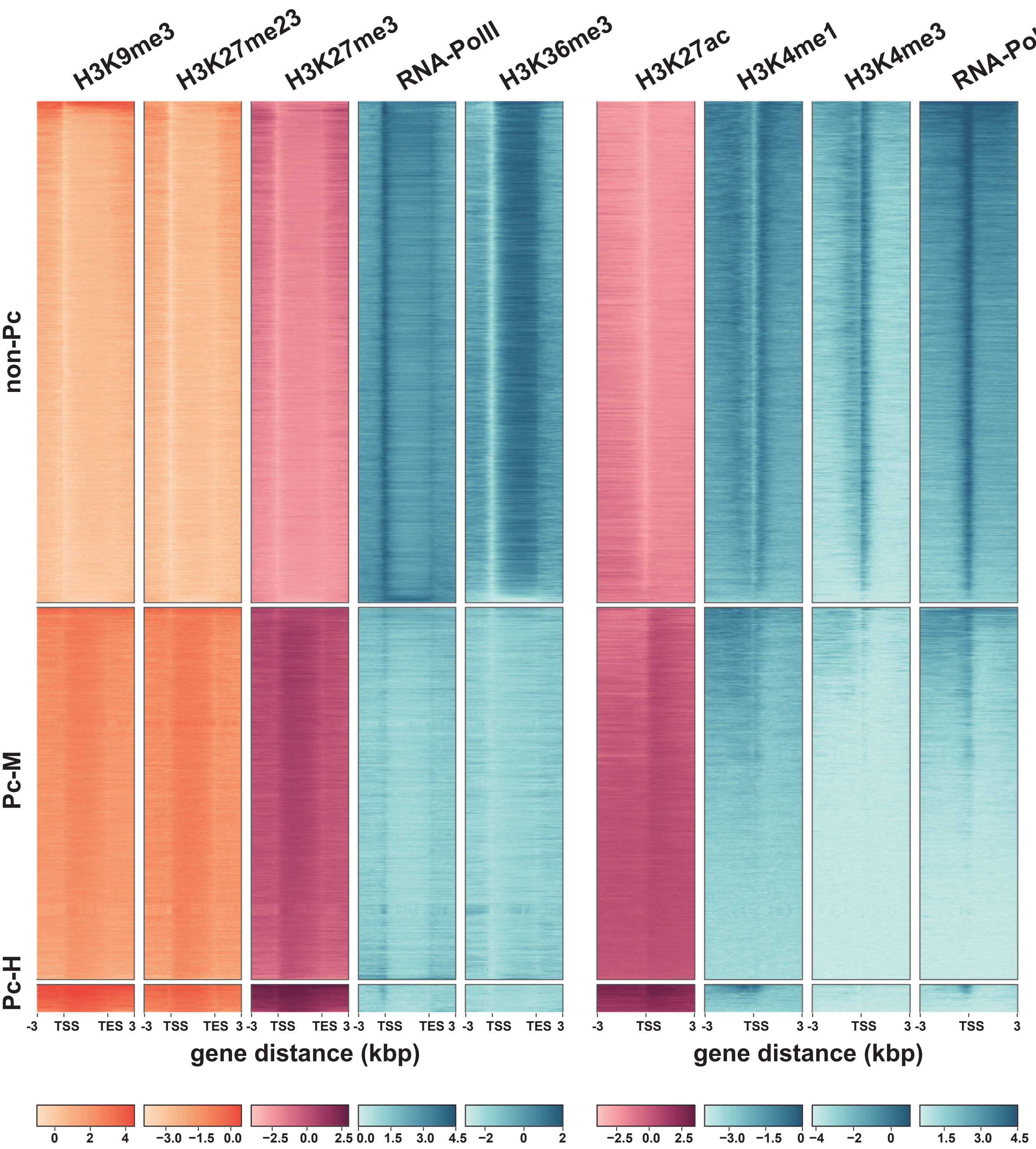

H3K9me3
H3K27me23
H3K27me3
RNA-PolII
H3K36me3
H3K27ac
H3K4me1
H3K4me3
RNA-PolII
non-Pc
Pc-M
Pc-H
-3 TSS TES 3
-3 TSS 3
gene distance (kbp)
gene distance (kbp)
0 2 4
−3.0 −1.5 0.0
−2.5 0.0 2.5
0.0 1.5 3.0 4.5
−2 0 2
−2.5 0.0 2.5
−3.0 −1.5 0
−4 −2 0
1.5 3.0 4.5

**Figure S8. Modification profiles for ClassFinder gene sets.**

Heatmaps of ChIP-seq signals for individual genes that are shown as average signals in Figure 1F. H3K4me1 and H3K4me3 were mapped around the transcriptional start site (TSS); Pol II and H3K36me3 along the gene body using Deeptools. The resulting heatmaps were split into groups according to gene states identified by ClassFinder (non-Pc, Pc-M, Pc-H) and the color scales below the heatmaps represent signal as log2 ratio over Input.

### Figure S9

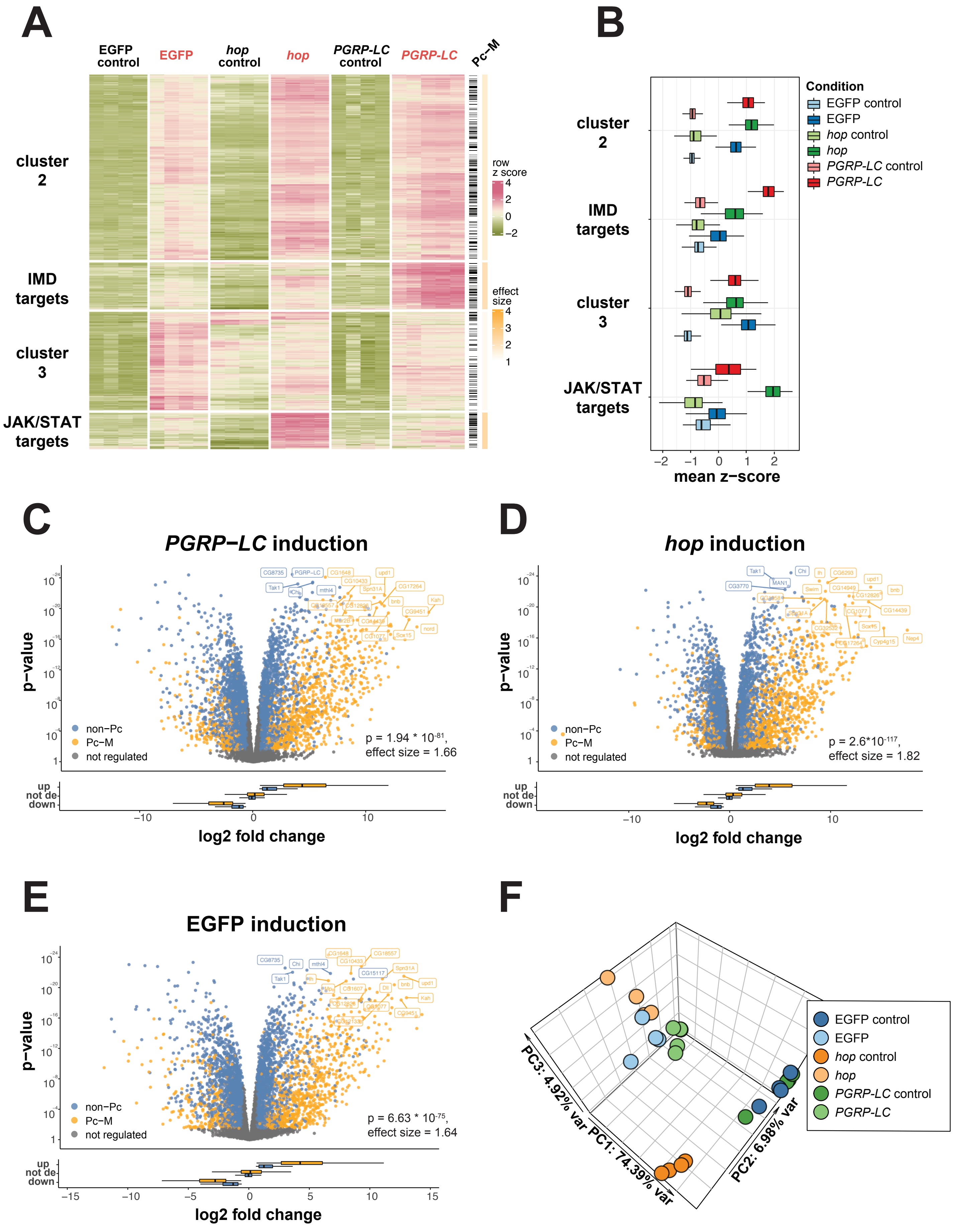

A
EGFP control
EGFP
hop control
hop
PGRP-LC control
PGRP-LC
Pc-M
cluster 2
IMD targets
cluster 3
JAK/STAT targets
row z score
effect size
B
Condition
EGFP control
EGFP
hop control
hop
PGRP-LC control
PGRP-LC
mean z-score
C
PGRP-LC induction
p-value
non-Pc
Pc-M
not regulated
p = 1.94 * 10^-81, effect size = 1.66
log2 fold change
D
hop induction
p = 2.6*10^-117, effect size = 1.82
E
EGFP induction
p = 6.63 * 10^-75, effect size = 1.64
F
PC1: 74.39% var
PC2: 6.98% var
PC3: 4.92% var

**Figure S9. Pc-M genes are induced by ectopic activation of signalling pathways.**

(**A**) Heatmap of genes highly up-regulated upon mifepristone induced expression of EGFP, *hopscotch* (*hop*) or *PGRP-LC* relative to non-induced controls. RNA-seq data was normalized using DESeq2 rlog and z-score (color scale: row z-score). We selected genes that were highly up-regulated (lfc >= 1 and p <= 0.00001) in at least one induction and clustered them by k-means clustering into 4 groups. For each cluster, we calculated the enrichment of Pc-M genes (right, black marks) over background using a hypergeometric test (color scale: effect size). IMD and JAK-STAT pathway dependent clusters were highly enriched for Pc-M genes. **IMD**: $p < 10^{-14}$, effect size = 2.1. **JAK-STAT**: $p < 10^{-15}$, effect size = 2.3. In addition, we found two clusters of genes that were induced either in response to mifepristone or as a non-specific result of GAL4 activation (cluster 2, 3). These clusters were also enriched for Pc-M genes, which is consistent with the notion that Pc-M genes are inducible. (**B**) Mean z-score of clusters in (A) across conditions. (**C-E**) Volcano plots of differentially expressed genes after GAL4 activation (mifepristone treatment vs. no treatment) for the induction of *PGRP-LC* (C), *hop* (D) and EGFP (E) expression. Volcanos show log2 fold change against p-value for differentially expressed non-Pc (blue) and Pc-M (yellow) genes, as well as for genes not significantly regulated (grey). Annotated p-values and effect sizes refer to enrichment of Pc-M genes by hypergeometric test with false discovery rate correction. Lower panels show box-plots of log2 fold changes for non-Pc (blue) and Pc-M (yellow) genes that were up-regulated (up), not differentially expressed (not de) or down-regulated (down). (**F**) Principal Component Analysis (PCA) of RNA-seq samples. Gene counts were normalized using DESeq2 rlog and PCA was performed using 1000 genes with the highest expression variance.

### Figure S10

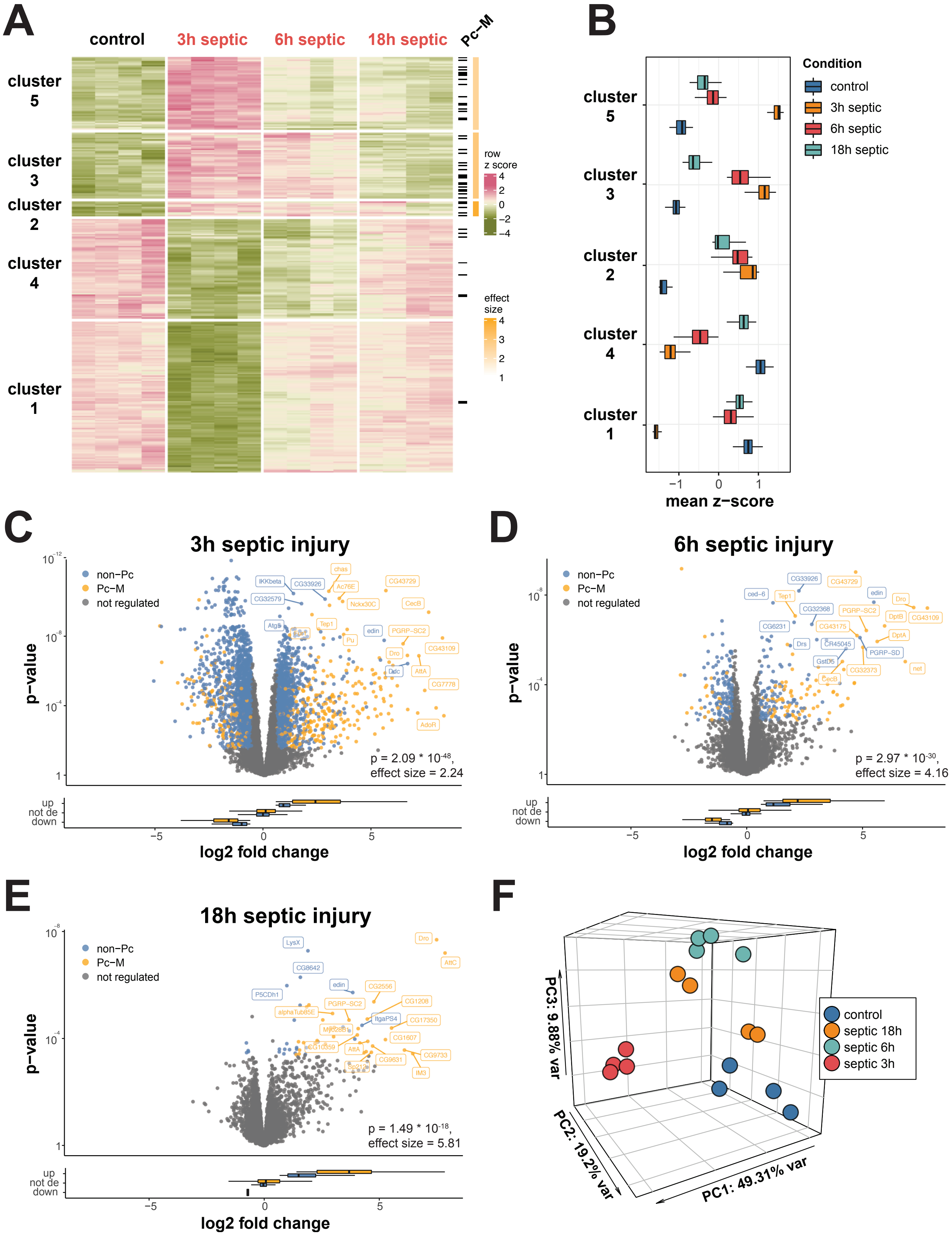

A
control
3h septic
6h septic
18h septic
Pc–M
cluster 5
cluster 3
cluster 2
cluster 4
cluster 1
row z score
4
2
0
−2
−4
effect size
4
3
2
1
B
Condition
control
3h septic
6h septic
18h septic
cluster 5
cluster 3
cluster 2
cluster 4
cluster 1
−1
0
1
mean z−score
C
3h septic injury
non−Pc
Pc−M
not regulated
p−value
p = 2.09 * 10-48,
effect size = 2.24
up
not de
down
−5
0
5
log2 fold change
D
6h septic injury
non−Pc
Pc−M
not regulated
p−value
p = 2.97 * 10-30,
effect size = 4.16
up
not de
down
−5
0
5
log2 fold change
E
18h septic injury
non−Pc
Pc−M
not regulated
p−value
p = 1.49 * 10-18,
effect size = 5.81
up
not de
down
−5
0
5
log2 fold change
F
PC3: 9.88% var
PC2: 19.2% var
PC1: 49.31% var
control
septic 18h
septic 6h
septic 3h

**Figure S10. Pc-M genes are induced in a septic injury infection model.**

(**A**) Heatmap of genes highly up-regulated in plasmatocytes after septic injury. RNA-seq data from larval plasmatocytes at different time points after septic injury were normalized using DESeq2 rlog and z-score (color scale: row z-score). We selected 300 genes with highest mean negative log p-value across all comparisons and clustered them by k-means clustering into 5 groups. For each cluster, we calculated the enrichment of Pc-M genes (right, black marks) over background using a hypergeometric test (color scale: effect size). All clusters of immune induced genes (cluster 2, 3 and 5) were highly enriched for Pc-M genes. **Cluster 2**: p = $6.9*10^{-4}$, effect size = 3.6. **Cluster 3**: p = $2.5*10^{-4}$, effect size = 2.8. **Cluster 5**: p = $1.9*10^{-3}$, effect size = 2.4. (**B**) Mean z-score of clusters in (A) across conditions. (**C-E**) Volcano plots of differentially expressed genes 3 h post septic injury (C), 6 h post septic injury (D) and 18 h post septic injury (E) (plasmatocytes from infected vs. uninfected larvae). Volcanos show log2 fold change against p-value for differentially expressed non-Pc (blue) and Pc-M (yellow) genes, as well as for genes not significantly regulated (grey). Annotated p-values and effect sizes refer to enrichment of Pc-M genes by hypergeometric test with false discovery rate correction. Lower panels show box-plots of log2 fold changes for non-Pc (blue) and Pc-M (yellow) genes that were up-regulated (up), not differentially expressed (not de) or down-regulated (down). (**F**) Principal Component Analysis (PCA) of RNA-seq samples. Gene counts were normalized using DESeq2 rlog and PCA was performed using 1000 genes with the highest expression variance.

### Figure S11

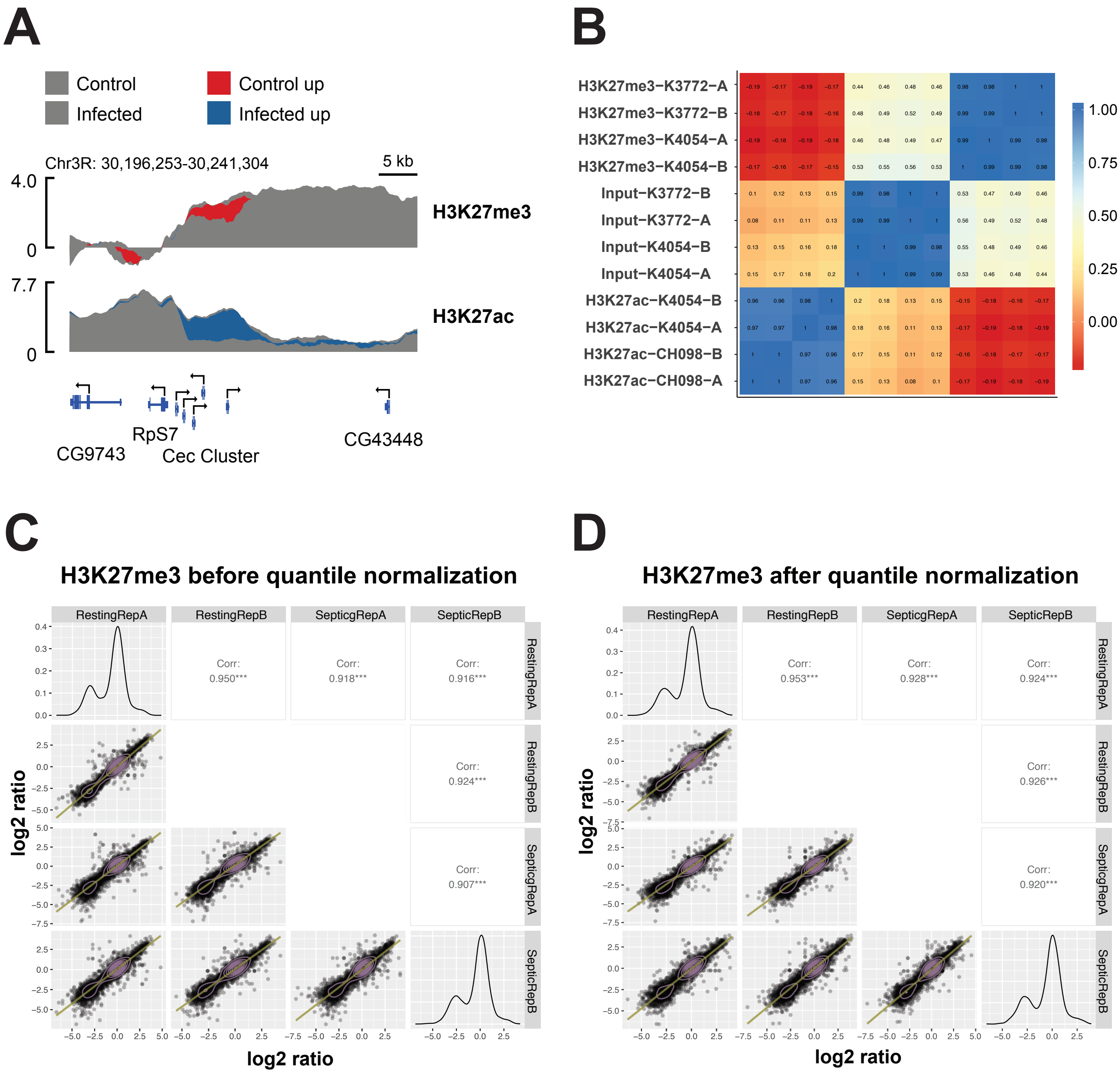
A
Control
Infected
Control up
Infected up
Chr3R: 30,196,253-30,241,304
5 kb
H3K27me3
H3K27ac
CG9743
RpS7
Cec Cluster
CG43448
B
H3K27me3−K3772−A
H3K27me3−K3772−B
H3K27me3−K4054−A
H3K27me3−K4054−B
Input−K3772−B
Input−K3772−A
Input−K4054−B
Input−K4054−A
H3K27ac−K4054−B
H3K27ac−K4054−A
H3K27ac−CH098−B
H3K27ac−CH098−A
C
H3K27me3 before quantile normalization
RestingRepA
RestingRepB
SepticgRepA
SepticRepB
Corr: 0.950***
Corr: 0.918***
Corr: 0.916***
Corr: 0.924***
Corr: 0.907***
log2 ratio
D
H3K27me3 after quantile normalization
Corr: 0.953***
Corr: 0.928***
Corr: 0.924***
Corr: 0.926***
Corr: 0.920***

**Figure S11. ChIP-seq of plasmatocytes at 6 h after septic injury.**

(**A**) Genome track of the genomic region surrounding the Cecropin (Cec) gene cluster, which are genes coding for immune induced antimicrobial peptides. ChIP-seq log2 ratio signals were plotted using SparK. Genes are marked at the bottom and ChIP targets on the right. Red regions indicate increased signal in non-infected plasmatocytes, while blue indicates higher signal after septic injury. (**B**) Heatmap of sample correlation. Mapped reads from each individual ChIP-seq replicate were binned using 10 kb windows and pairwise Spearman correlation coefficients calculated using Deeptools. Samples marked by K3772 and CH098 are from uninfected plasmatocytes, while K4054 samples are post septic injury. All samples from the same ChIP targets are highly correlated (color scale indicates correlation coefficients). (**C-D**) H3K27me3 signal before (C) and after (D) quantile normalization. Gene-level read counts are plotted as pairwise correlations, with trend-lines in brown. Before normalization (C) some point clouds are "banana-shaped" (e.g. RestingRepB vs SepticRepA) and trendlines deviate from the ideal x = y diagonal at the extremes. After normalization (D) the point clouds are more homogeneous, and the trendlines are closer to the diagonal.

### Figure S12

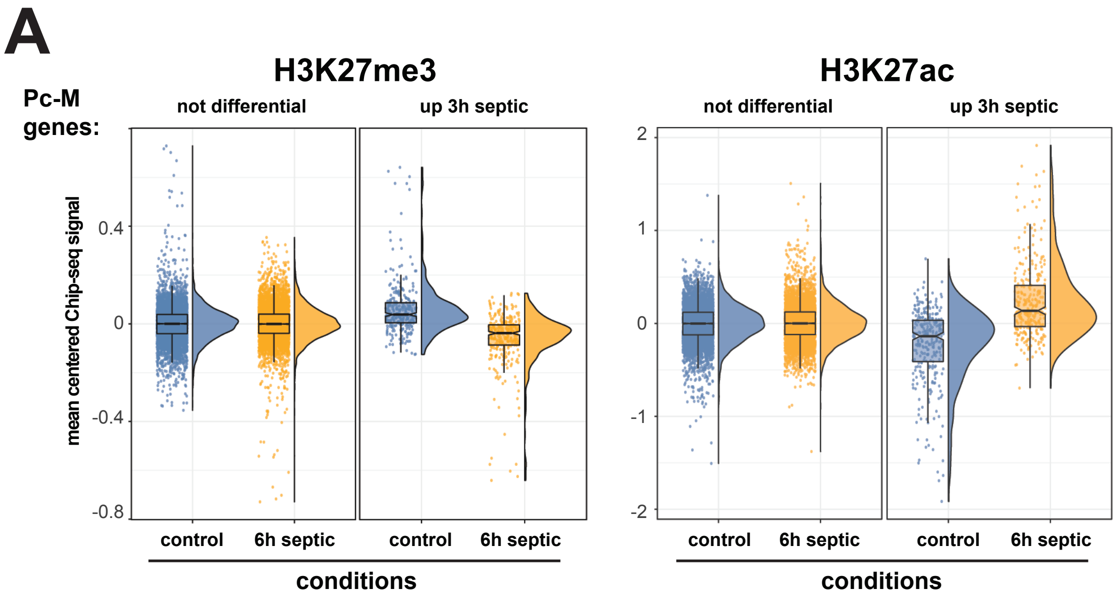
A
H3K27me3
H3K27ac
Pc-M
genes:
not differential
up 3h septic
not differential
up 3h septic
mean centered Chip-seq signal
0.4
0
-0.4
-0.8
2
1
0
-1
-2
control
6h septic
control
6h septic
control
6h septic
control
6h septic
conditions
conditions

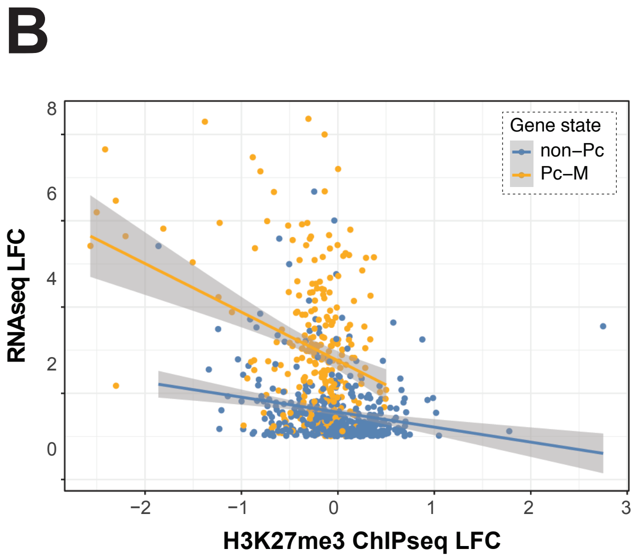
B
Gene state
non−Pc
Pc−M
RNAseq LFC
8
6
4
2
0
−2
−1
0
1
2
3
H3K27me3 ChIPseq LFC

**Figure S12. Immune induced genes loose H3K27me3 and gain H3K27ac after septic injury.**

(**A**) Differential modification of H3K27me3 and H3K27ac at Pc-M genes that were selected based on their differential expression at 3 h post septic injury. Pc-M genes were grouped into genes not differentially regulated at 3 h post septic injury (not differential) and up-regulated genes (up 3 h septic, $lfc > 0$, $p < 0.05$). For each gene in these groups mean centered ChIP-seq signals were determined in control plasmatocytes (blue) and in plasmatocytes 6 h after septic injury (yellow). These signals were derived from quantile normalized gene-level H3K27me3 or H3K27ac log2 ratio read counts as mean from both replicates. For up-regulated genes (up 3 h septic) H3K27me3 levels (Mann-Whitney-U $p = 7.93*10^{-24}$) and H3K27ac levels ($p = 6.17*10^{-20}$) were significantly different. (**B**) Correlation between loss of H3K27me3 and RNA-seq based gene induction by after septic injury. For all genes up-regulated at 3 h post septic injury, ChIP-seq mean log2 fold change (from quantile normalized data) was plotted against RNA-seq log2 fold change. Pc-M genes are marked in yellow and non-Pc genes in blue. Lines show linear models with 95% confidence interval in grey. For Pc-M there is a statistically significant correlation ($r = -0.32$, $p = 1.44*10^{-8}$).

### Figure S13

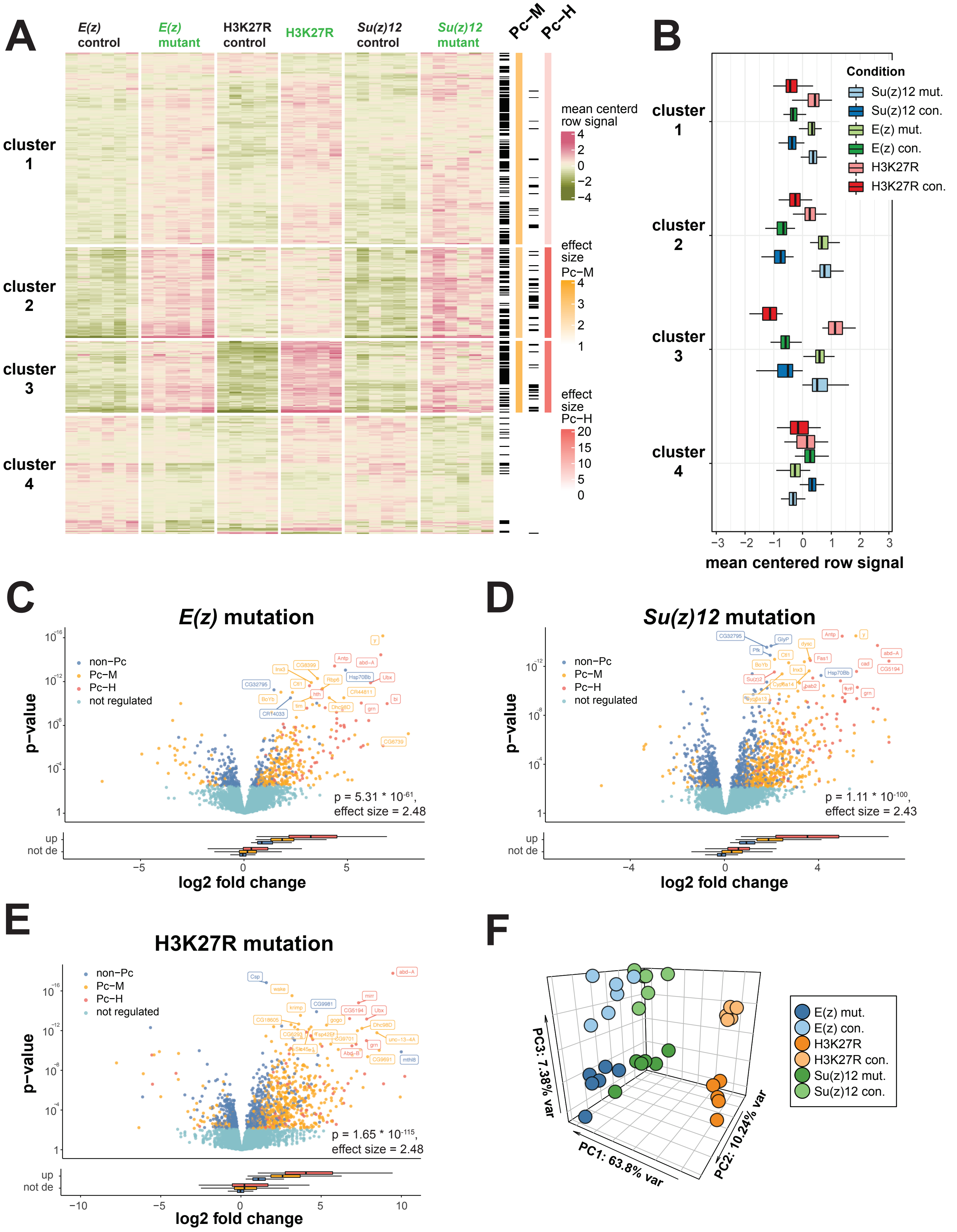

A
E(z) control
E(z) mutant
H3K27R control
H3K27R
Su(z)12 control
Su(z)12 mutant
Pc−M
Pc−H
cluster 1
cluster 2
cluster 3
cluster 4
mean centerd row signal
effect size Pc−M
effect size Pc−H
B
Condition
Su(z)12 mut.
Su(z)12 con.
E(z) mut.
E(z) con.
H3K27R
H3K27R con.
mean centered row signal
C
E(z) mutation
non−Pc
Pc−M
Pc−H
not regulated
p−value
p = 5.31 * 10^-61, effect size = 2.48
up
not de
log2 fold change
D
Su(z)12 mutation
p = 1.11 * 10^-100, effect size = 2.43
E
H3K27R mutation
p = 1.65 * 10^-115, effect size = 2.48
F
E(z) mut.
E(z) con.
H3K27R
H3K27R con.
Su(z)12 mut.
Su(z)12 con.
PC1: 63.8% var
PC2: 10.24% var
PC3: 7.38% var

**Figure S13. PRC2 and H3K27me3 repress the expression of Pc-M and Pc-H genes.**
(**A**) Heatmap of genes highly regulated in plasmatocytes after removing PRC2 activity (*E(z)* mutant, *Su(z)12* mutant) or H3K27me3 (H3K27R) relative to non-mutant control cells (control). RNA-seq data were normalized using DESeq2 rlog and z-score (color scale: row z-score). We selected 500 genes with highest mean negative log p-value across all comparisons and clustered them by k-means clustering into 4 groups. For each cluster, we calculated the enrichment of Pc-M and Pc-H genes (right, black marks) over background using a hypergeometric test (yellow color scale: effect size Pc-M, red color scale: effect size Pc-H). All clusters of genes upregulated by loss of H3K27me3 or PRC2 (cluster 1, 3 and 4) are highly enriched for Pc-M and Pc-H genes. **Cluster 1**: Pc-M, $p < 10^{-19}$, effect size = 3.4; Pc-H, $p < 10^{-18}$, effect size = 18. **Cluster 3**: Pc-M, $p < 10^{-14}$, effect size = 2.8; Pc-H, p < 10E-27, effect size = 20. **Cluster 4**: Pc-M, p < 10E-40, effect size = 3; Pc-H, $p < 10^{-19}$, effect size = 6. (**B**) Mean centered gene signal of clusters in (A) across conditions. (**C-E**) Volcano plots of differentially expressed genes after loss of PRC2 activity or H3K27me3 (mutant plasmatocytes vs control cells) by *E(z)* (C), *Su(z)12* (D) or H3K27R (E) mutations. Volcanos show log2 fold change against p-value for differentially expressed non-Pc (blue), Pc-M (yellow) and Pc-H (red) genes, as well as for genes not significantly regulated (turquoise). Annotated p-values and effect sizes refer to enrichment of Pc-M genes by hypergeometric test with false discovery rate correction. Lower panels show box-plots of log2 fold changes for non-Pc (blue) Pc-M (yellow) and Pc-H (red) genes that were up-regulated (up), not differentially expressed (not de). (**F**) Principal Component Analysis (PCA) of RNA-seq samples. Gene counts were normalized using DESeq2 rlog and PCA was performed using 1000 genes with the highest expression variance.

### Figure S14

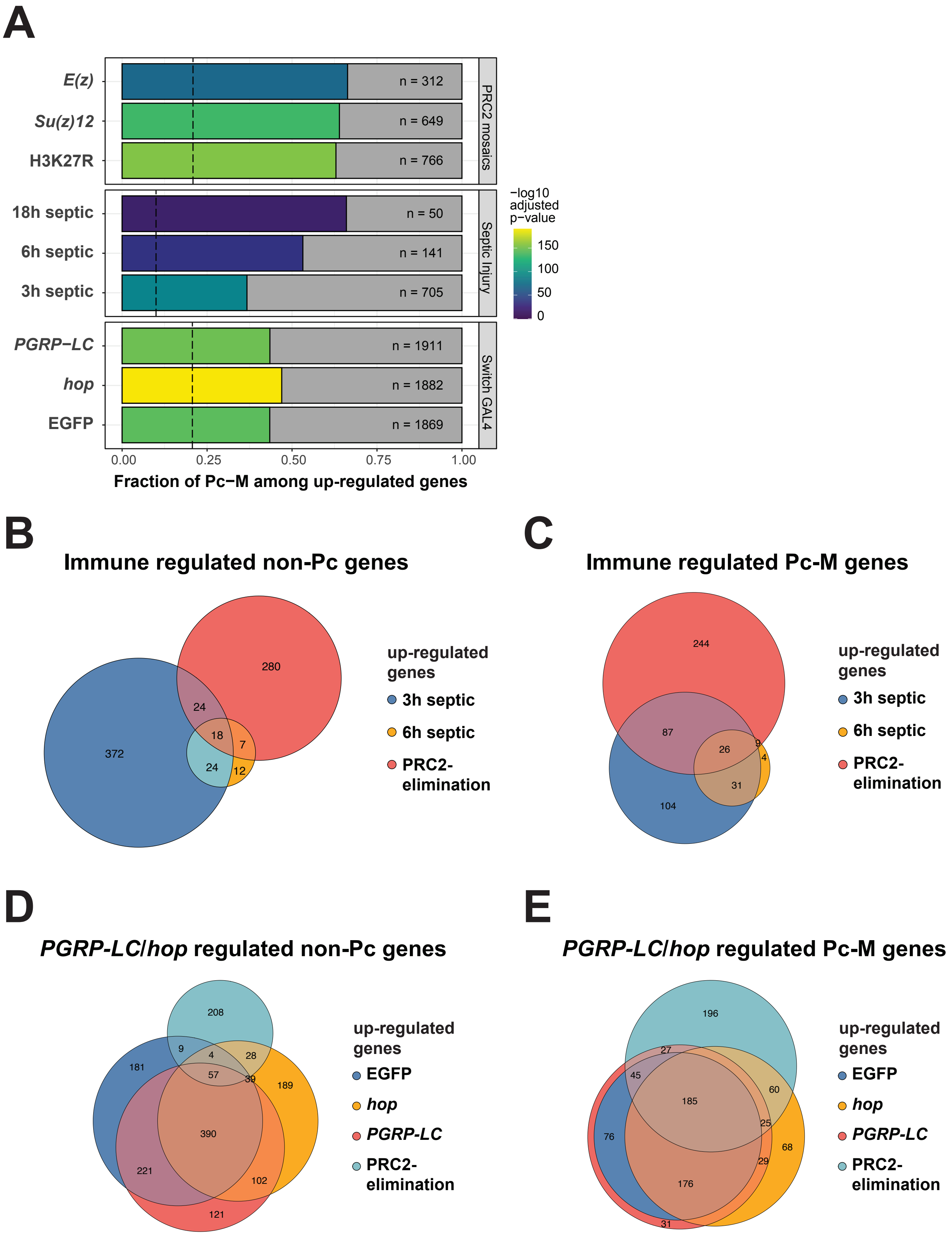
A
E(z)
Su(z)12
H3K27R
18h septic
6h septic
3h septic
PGRP−LC
hop
EGFP
n = 312
n = 649
n = 766
n = 50
n = 141
n = 705
n = 1911
n = 1882
n = 1869
PRC2 mosaics
Septic Injury
Switch GAL4
−log10 adjusted p−value
150
100
50
0
0.00
0.25
0.50
0.75
1.00
Fraction of Pc−M among up-regulated genes
B
Immune regulated non-Pc genes
280
24
18
7
372
24
12
up-regulated genes
3h septic
6h septic
PRC2-elimination
C
Immune regulated Pc-M genes
244
87
26
9
4
31
104
up-regulated genes
3h septic
6h septic
PRC2-elimination
D
PGRP-LC/hop regulated non-Pc genes
208
9
4
28
181
57
39
189
390
221
102
121
up-regulated genes
EGFP
hop
PGRP-LC
PRC2-elimination
E
PGRP-LC/hop regulated Pc-M genes
196
27
45
60
185
25
76
68
29
176
31
up-regulated genes
EGFP
hop
PGRP-LC
PRC2-elimination

**Figure S14. Inducible Pc-M genes are repressed against transcriptional activation by PRC2 and H3K27me3.**

(**A**) Enrichment of Pc-M genes among up-regulated genes after PRC2 elimination (top), septic injury (middle) and mifepristone induced expression (bottom). From the respective comparisons of RNA-seq datasets, up-regulated genes were identified (lfc > 0, $p < 0.05$) and enrichment of Pc-M genes over background (all genes analyzed in the respective dataset) was determined by a hypergeometric test. Bar fill represents the p-value, bar length the fraction of Pc-M genes, the dotted line represents the expected fraction under the null hypothesis and n indicates the total number of up-regulated genes. (**B-E**) Venn diagrams showing overlap of genes up-regulated in any of the conditions eliminating PRC2 function (combined *E(z)*, *Su(z)12* or H3K27R data sets) with genes that were induced by 3 h and 6 h after septic injury (B,C) or in response to mifepristone induced expression of EGFP, *hopscotch* (*hop*) or *PGRP-LC* (D,E).

### Figure S15

## A

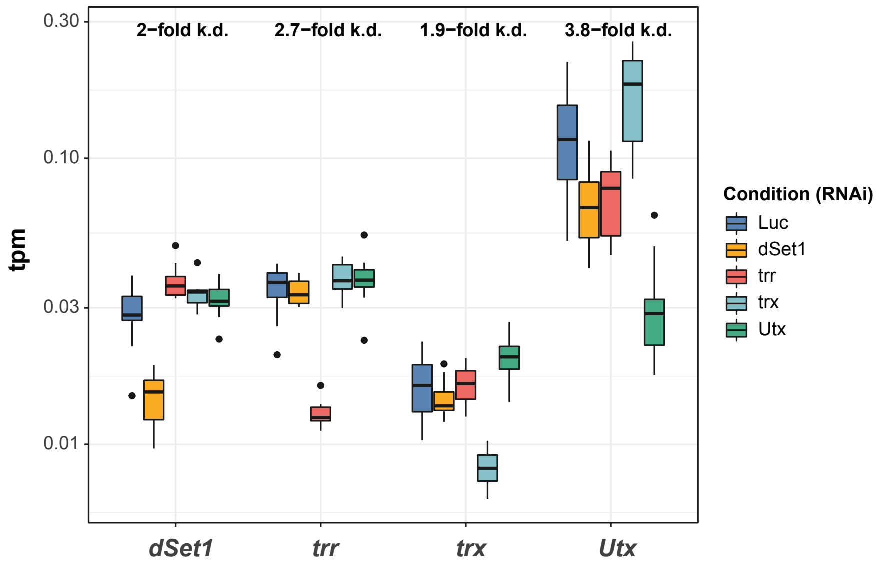

## B

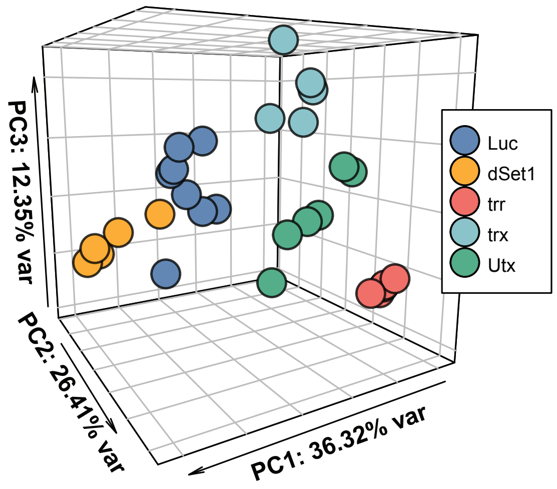

**Figure S15. RNAi mediated knockdown of TrxG proteins in plasmatocytes.**

(**A**) Knockdown efficiencies in plasmatocytes. Expression of *dSet1*, *trr*, *trx* and *Utx* was determined by tags per million (tpm) in knockdown plasmatocytes. Knockdowns were induced by expression of RNAi transgenes targeting *dSet1*, *trr*, *trx* and *Utx* and Luciferase (Luc) as a control. Knockdown efficiencies for the constructs (relative to Luciferase control) are marked at the top. (**B**) Principal Component Analysis (PCA) of RNA-seq from *dSet1*, *trr*, *trx*, *Utx* and Luciferase RNAi-knockdown plasmatocytes. Gene counts were normalized using DESeq2 rlog and PCA was performed using 1000 genes with the highest expression variance.

### Figure S16

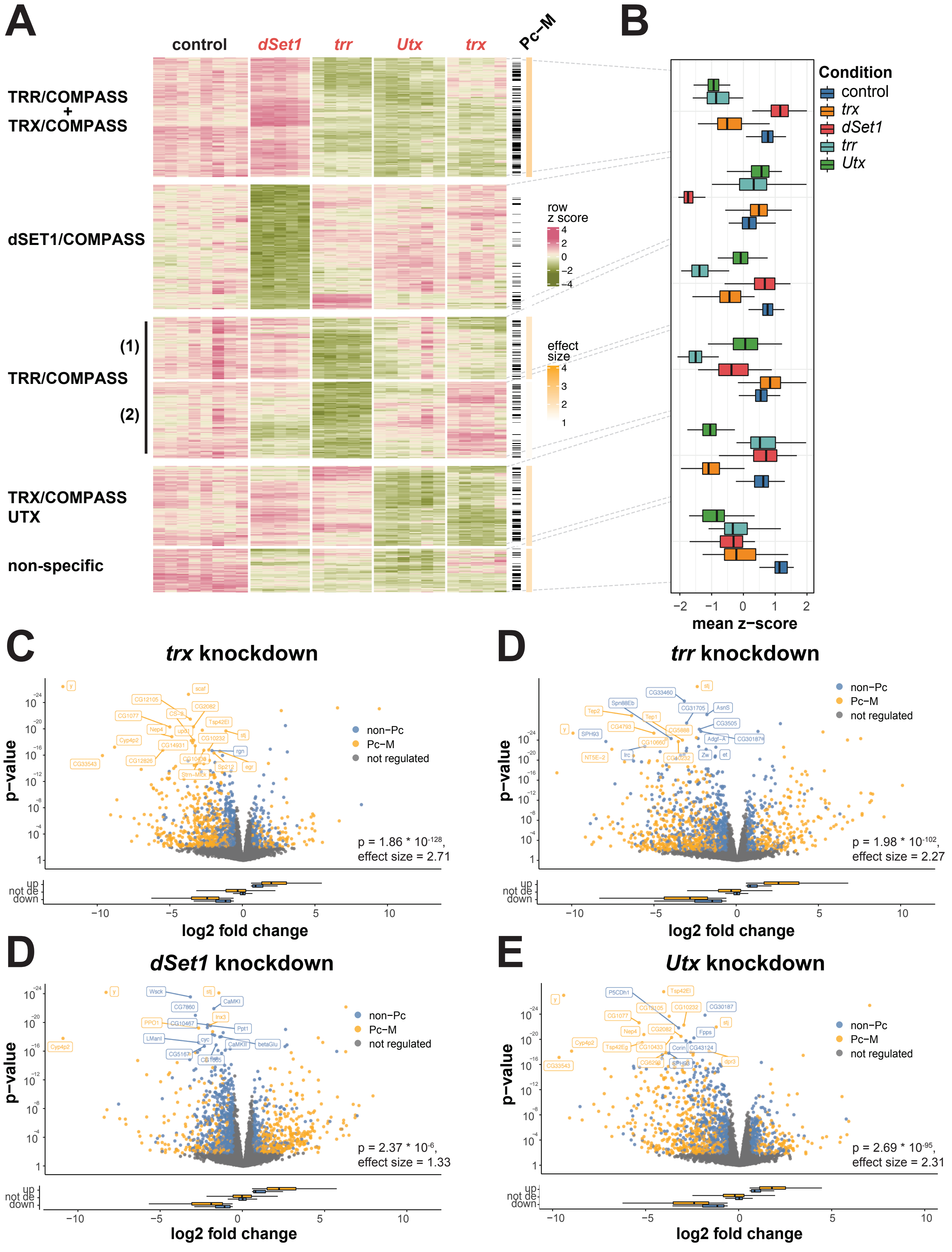
A
control
dSet1
trr
Utx
trx
Pc−M
TRR/COMPASS
+
TRX/COMPASS
dSET1/COMPASS
(1)
TRR/COMPASS
(2)
TRX/COMPASS
UTX
non-specific
row
z score
4
2
0
−2
−4
effect
size
4
3
2
1
B
Condition
control
trx
dSet1
trr
Utx
−2
−1
0
1
2
mean z−score
C
trx knockdown
p−value
non−Pc
Pc−M
not regulated
p = 1.86 * 10^-128,
effect size = 2.71
up
not de
down
log2 fold change
D
trr knockdown
p = 1.98 * 10^-102,
effect size = 2.27
D
dSet1 knockdown
p = 2.37 * 10^-6,
effect size = 1.33
E
Utx knockdown
p = 2.69 * 10^-95,
effect size = 2.31

**Figure S16. TrxG-proteins positively regulate homeostatic expression of Pc-M genes**

(**A**) Heatmap of genes highly down-regulated upon RNAi mediated knockdown of *dSet1*, *trr*, *Utx* and *trx* relative to control knockdown of Luciferase in plasmatocytes. RNA-seq data was normalized using DESeq2 rlog and z-score (color scale: row z-score). We selected the top 1000 genes with the highest mean negative log p-value among down-regulated genes across all comparisons and clustered them by k-means clustering into 6 groups. For each cluster, we calculated the enrichment of Pc-M genes (right, black marks) over background using a hypergeometric test (color scale: effect size). Clustering identified genes influenced by TRX/COMPASS (*trx* knockdown), TRR/COMPASS (*trr* and *Utx* knockdown), dSET1/COMPASS (*dSet1* knockdown), TRX/COMPASS and UTX (*trx* and *Utx* knockdowns) and no specific COMPASS complex (*trx*, *trr*, *Utx* and *dSet1* knockdowns). Several clusters were significantly enriched for Pc-M genes. **TRX/COMPASS and TRR/COMPASS**: $p < 10^{-25}$, effect size = 2.5. **TRR/COMPASS (1)**: $p = 1.3*10^{-6}$, effect size = 1.9). **TRX/COMPASS and UTX**: $p = 2.2*10^{-4}$, effect size = 1.6. **Non-specific**: $p = 4.4*10^{-7}$, effect size = 2.1. (**B**) Mean z-score of clusters in (A) across conditions. (**C-F**) Volcano plots of differentially expressed genes after knockdown of *trx* (C), *trr* (D), *dSet1* (E) and *Utx* (F) (knockdown plasmatocytes vs control cells). Volcanos show log2 fold change against p-value for differentially expressed non-Pc (blue) and Pc-M (yellow) genes, as well as for genes not significantly regulated (grey). Annotated p-values and effect sizes refer to enrichment of Pc-M genes by hypergeometric test with false discovery rate correction. Lower panels show box-plots of log2 fold changes for non-Pc (blue) and Pc-M (yellow) genes that were up-regulated (up), not differentially expressed (not de) or down-regulated (down).

### Figure S17

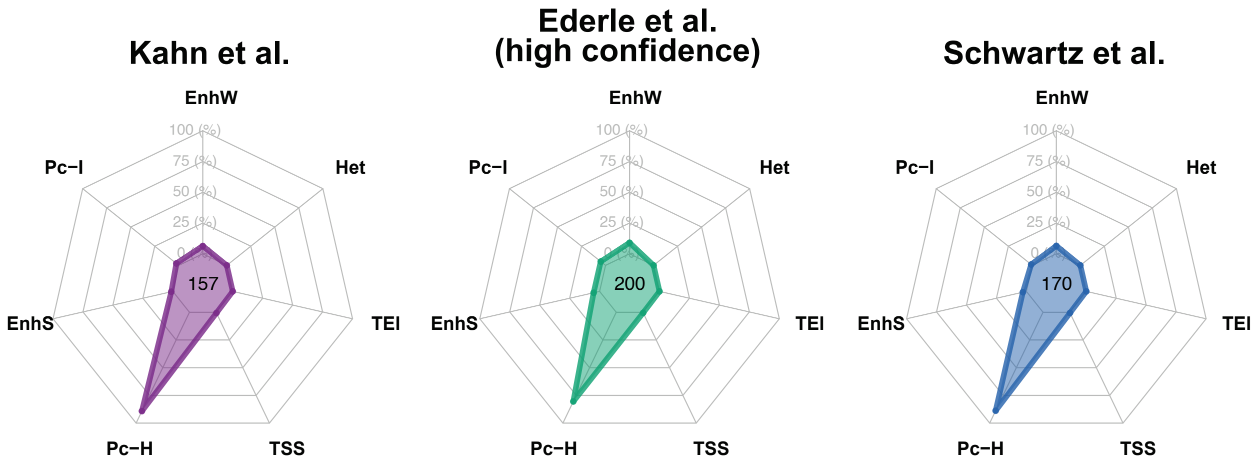
Kahn et al.
Ederle et al.
(high confidence)
Schwartz et al.
EnhW
Het
TEI
TSS
Pc−H
EnhS
Pc−I
100 (%)
75 (%)
50 (%)
25 (%)
0 (%)
157
200
170

**Figure S16. Well studied PREs are localized to Pc-H chromatin.**

Annotated Polycomb response elements (PREs) were identified from datasets of Kahn *et al.* (*54*), Ederle *et al.* (55) (high confidence set only) and Schwartz *et al*. (56). From these sets we identified overlap with the different chromatin states from the 7-state ClassFinder model. Relative frequency of overlap is shown as a radar plot, with the number of PREs in the corresponding dataset in the middle. PREs locate almost entirely to Pc-H chromatin across all 3 reference data sets.
